# MPK3 mediated phosphorylation of WRKY48 down regulates CIPK6 expression during *Pst*DC3000 challenge in Arabidopsis

**DOI:** 10.1101/2025.05.21.655431

**Authors:** Nidhi Singh, Debasis Chattopadhyay

## Abstract

WRKY transcription factor (TF) family are one of the key regulators of plant immune responses, functioning as both activators and suppressors depending on the pathogen encountered. Among these, WRKY48, a class IIC TF, functions as a negative regulator of pattern-triggered immunity (PTI) in *Arabidopsis thaliana*. Our study explicates the regulatory mechanisms underlying the function of WRKY48, revealing its role in repressing the expression of *CIPK6*, a negative regulator of immune response in *Arabidopsis*, during *Pseudomonas syringae* DC3000 (*Pst*DC3000) infection. This suppression is mediated through MPK3-dependent phosphorylation of WRKY48*. In vitro* phosphorylation assays demonstrated that MPK3 primarily phosphorylates WRKY48 at serine residues S^35^ and S^40^. The phosphorylated WRKY48 predominantly localizes to the nucleus, enhancing its DNA-binding affinity and trans-repressive activity at specific W-box motifs in the CIPK6 promoter. Whereas invivo dual luciferase shows that unphosphorylated WRKY48 (S^26^S^35^S^40^ / A^26^A^35^A^40^ ) unable to repress luciferase activity of *CIPK6* promoter. The bioinformatics analysis identified two WRKY-binding sites (P1 and P2) on the CIPK6 promoter and electrophoretic mobility shift assays confirms the preferential binding of WRKY48 to the P1 site. Unlike other WRKY TFs (WRKY28 and WRKY8) in the same subgroup, WRKY48 exhibited unique specificity, highlighting the distinct regulatory roles within the WRKY family. Functional studies demonstrated that phosphorylation of WRKY48 enhances its ability to suppress CIPK6 both transcriptionally and translationally, leading to increased reactive oxygen species (ROS) levels and decreased susceptibility to *Pst*DC3000. The reduced expressions of PTI marker genes and ROS-associated genes in *wrky48* mutants were associated with enhanced resistance in *wrky48* and *cipk6* mutants and emphasizes the critical role of WRKY48 in fine-tuning immune responses. Our findings identify a novel MPK3-WRKY48-CIPK6 signaling module that connects calcium-dependent pathways to MAPK signalling to modify plant immunity.

**Highlights:**

- WRKY48, a class IIC transcription factor, functions as a negative regulator of pattern-triggered immunity (PTI) in *Arabidopsis thaliana* during Pseudomonas syringae DC3000 (PstDC3000) infection.
- WRKY48 suppresses the expression of CIPK6, a negative regulator of immune signaling, in response to bacterial infection.
- MAP kinase MPK3 phosphorylates WRKY48 at serine residues S^35^ and S^40^, enhancing its nuclear localization and transcriptional repressive activity
- Phosphorylated WRKY48 shows increased binding affinity to W-box motifs in the *CIPK6* promoter, especially at the P1 binding site, as confirmed by electrophoretic mobility shift assays (EMSA).
- Unlike WRKY28 and WRKY8, WRKY48 exhibits unique binding specificity, suggesting functional diversification within WRKY subgroup IIC.
- Suppression of CIPK6 by phosphorylated WRKY48 leads to: Elevated ROS accumulation, Reduced PTI marker gene expression, Increased susceptibility to *PstDC3000*

## Introduction

Plants activate their innate immune system in response to threats from pathogens (Bentham et al., 2020). Pathogens are recognised through plant surface receptors i.e. pattern recognition receptors (PRRs) that results in the activation of pattern triggered immunity (PTI) (Yuan et al., 2021). The successful pathogen releases specific effector molecules into the plant cells. Those effectors are recognised by the nuclear-leucine-repeat rich receptors (NLRs) and activate effector triggered immunity (ETI), that suppresses PTI (Thordal-Christensen, 2020; Ngou et al., 2022; Waheed et al., 2022). Defense responses in both PTI and ETI are mediated by hormone signaling pathways that vary depending on the type of pathogen. Key phytohormones, such as salicylic acid (SA), jasmonic acid (JA), ethylene (ET), and abscisic acid (ABA), play differential roles ( Bürger and Chory, 2019; Aerts et al., 2021) in immune response. Upon pathogen recognition, various regulatory elements orchestrate transcriptional reprogramming to activate defense responses (Tsuda and Somssich, 2015). This transcriptional reprogramming involves transcription factors (TFs), chromatin remodelers, and RNA/DNA modulators, which modify the transcriptome and integrate signaling pathways to generate diverse transcriptional outcomes (Moore et al., 2011)(Tsuda and Somssich, 2015; Li et al., 2016). These molecular regulators often undergo post-translational modifications, including phosphorylation, sumoylation, and ubiquitination, enabling precise modulation of immune responses. Pathogens exploit these modifications, targeting key regulators like MAPK signaling pathways to facilitate infection (Newton et al., 2007; Ribet and Cossart, 2010; Gorczyca et al., 2024).

The effective transcriptional control of defense genes is critical, with TFs playing a central role. Transcription factors such as WRKYs, ERFs (Ethylene Responsive factors), bZIPs (Basic Leucine Zipper Domains), bHLHs (basic Helix Loop helix), MYBs (v-Myb myeloblastosis viral oncogene homolog), CAMTAs (Calmodulin Transcription Activators), and CBP60s (Calmodulin-binding protein 60)/SARD1 (SYSTEMIC ACQUIRED RESISTANCE DEFICIENT 1) are known to be involved in developing resistance against pathogens (Fernández-Calvo et al., 2011; Phukan et al., 2016; Kim et al., 2020; Wang et al., 2023). WRKY TFs are particularly versatile, influencing biotic and abiotic stress responses, as well as developmental processes (Jiang et al., 2015). They often form homo-and heterocomplexes to regulate gene expression via auto-and cross-regulatory mechanisms by interacting with various other proteins, including receptors, kinases, and other TFs, forming intricate transcriptional regulatory networks. WRKY TFs typically bind W-box (TTGACY, core sequence TGAC) cis-elements in gene promoters but can also recognize non-W-box element (Bakshi and Oelmüller, 2014).

WRKY proteins are categorised into three classes according to the presence of WRKY domains (WDs) and zinc finger motifs. Classes I and II contain C2H2 zinc fingers, while group III features C2HC zinc fingers. Phylogenetic analyses further divide class II into at least five subclasses (a-e). WRKY TFs exhibit diverse roles in plant immunity. Loss-and gain-of-function studies in Arabidopsis reveal that WRKY TFs function as differential regulators in defense against biotrophic and necrotrophic pathogens (Eulgem et al., 2000; Eulgem and Somssich, 2007). For example, AtWRKY52, containing a TIR–NBS–LRR domain, works with RPS4 to defend against *Colletotrichum higginsianum* and *Pseudomonas* syringae (Narusaka et al., 2009). In another study, AtWRKY52 interacts with bacterial effector PopP2 secreted by Ralstonia solanacearum in the nucleus and plant become resistant (Deslandes et al., 2003 and Deslandes et al.,2002). Similarly, WRKY16 interact with RRS1 and TTR1 NBS-LRR proteins, and this interaction activates downstream defense related genes, indicating their involvement in ETI pathway (Chi et al., 2013). AtWRKY50/51 increases resistance to *Alternaria brassicicola* but increases susceptibility to *Botrytis cinerea* by suppressing JA signalling through the SA and low oleic acid pathways (Gao et al., 2011). Similarly, AtWRKY33, AtWRKY3, and AtWRKY4 also positively influence defense against fungal pathogen like *Alternaria brassicicola* and *Botrytis cinerea* (Zheng et al., 2006; Lai et al., 2008). A large number of WRKY transcription factors (TFs) function as negative regulators of defense sinaling. Mutants of AtWRKY7, AtWRKY11, and AtWRKY17 show increased susceptibility to *Pseudomonas syringae* (Journot-Catalino et al., 2006), while AtWRKY38 and AtWRKY62 (Mao et al., 2007) negatively affect basal resistance to this pathogen (Kim et al., 2008). Systemic acquired resistance (SAR) and basal resistance are adversely regulated by AtWRKY48 and -58, respectively (Wang et al., 2006). AtWRKY48 adversely impact basal resistance to *P. syringae*, with *atwrky48* mutants showing reduced bacterial growth and increased PR1 expression, while overexpression of AtWRKY48 makes plant susceptible to *Pseudomonas* (Xing et al., 2008).

Since WRKYs are considered as “Jack of All Trades” they also play peculiar roles. For example, AtWRKY27 mutation postpones symptoms of *Ralstonia solanacearum* (Mukhtar et al., 2008), while AtWRKY23 knockdown reduces vulnerability to cyst nematodes (Grunewald et al., 2008). While AtWRKY18, AtWRKY40, and AtWRKY60 provide variable resistance to other diseases like as powdery mildew and *Botrytis cinerea*, they redundantly reduce resistance to *P. syringae* (Xu et al., 2006). Because of their multiple roles, AtWRKY53 (Murray et al., 2007) and AtWRKY41 (Higashi et al., 2008) affect resistance to particular pathogens in different ways based on their expression or environment conditions. These results demonstrate the intricate functions of WRKY TFs in regulating plant immunity.

Early innate immunity, or pattern-triggered immunity (PTI), involves the parallel activation of MAP kinases (MPKs) (Meng and Zhang, 2013a) and Ca²⁺-dependent protein kinases (CPKs) (Singh et al., 2017). The mitogen-activated protein kinase (MPK) cascade is crucial in plant defense, with MPKs like MPK3, MPK4, and MPK6 activated via phosphorylation by MPK kinases (MKKs/MEKs) and upstream MPK kinase kinases (MEKKs) (Meng and Zhang, 2013a). A well-studied signaling pathway is flagellin-triggered basal defense, where MAPKs (3/4/6) are activated through flagellin perception via FLS2 receptor, followed by phosphorylation of MPK cascade. This signaling is rapid where MPK3/4/6 is activated within ∼5 minutes, peaking at ∼30 minutes (Mithoe and Menke, 2018). Whereas CPK-dependent signalling, which is initiated by Ca^2+^ influx through Ca^2+^ channels, peaks three minutes after elicitor (AtPeps) treatment and highest peak could be seen after one minute of Ca^2+^ accumulation (Qia et al., 2010).

A number of WRKY TFs are known to be substrate of MPKs that play key role in plant immunity. For example, AtWRKY33 is essential for AtMPK3/AtMPK6-mediated pathogen-induced camalexin and ethylene biosynthesis in Arabidopsis. It is phosphorylated by AtMPK3/AtMPK6 during *Botrytis cinerea* infection, with phosphorylation critical for its function in camalexin production (Mao et al., 2011). AtWRKY33 regulates its promoter in a positive-feedback loop and targets Phytoalexin Deficient3 (PAD3) for the final step of camalexin synthesis. It also interacts with the AtMPK4-AtMKS1 complex that is released upon *Pseudomonas syringae* challenge or flg22 treatment (Qui et al., 2008). Also, AtMPK3 destabilizes the WRKY46 during flg22 treatment and concurrently WRKY8, WRKY28, WRKY48, and CPKs (5, 7, and 11) interact with WRKY46 in Arabidopsis. CPKs phosphorylate the AtWRKY8, 28, and 48 and increase their binding to W-box elements. In contrast to MPKs, which are quickly activated by flagellin, CPKs are sustainedly triggered by avirulent effectors (Gao et al., 2013).

In this study, we report that WRKY48, a class IIC TFs, negatively impacts the generation of reactive oxygen species during activation of PTI. It regulates the expression of CBL-Interacting Protein kinase 6 (CIPK6) during *Pst*DC3000 challenge and flg22 treatment as well in Arabidopsis. AtWRKY48 primarily interacts with AtMPK3 in the nucleus. The phosphorylation of WRKY48 through AtMPK3 not only increases its binding to the W-box elements present on the CIPK6 promoter but also its trans-repression activity. Interestingly, WRKY48 phosphorylation is induced by MPK3 in response to the *Pst*DC3000 challenge that results in its enhanced trans-repression of the CIPK6 transcript *in planta*. Further, downregulation of CIPK6 results in resistance towards *Pst*DC3000 associated with enhanced *PR1* expression and ROS generation in WRKY48 overexpression plants as compared to *wrky48* mutants.

## Results

### WRKY48 binds to the *CIPK6* promoter

We have previously shown that CIPK6 acts as a negative regulator of immune response in Arabidopsis against *Pseudomonas* syringae pv. DC3000 (*Pst*) challenge. Upon infection by the bacteria, *CIPK6* transcript level was reduced quickly and restoration of *CIPK6* expression in the cipk6-/-line restored the phenotype of the wild type plants (Sardar et al., 2017). To investigate the transcriptional regulation of *CIPK6*, we analysed the 1.2 Kb-long upstream region of *CIPK6* transcriptional start site through PLANTPAN4.0 tool. We found various cis-acting elements suitable for binding of TFs including WRKY proteins. We searched for the WRKY proteins that are known to be involved in the negative regulation of plant immunity e.g. WRKY48 (Xing et al., 2008) and WRKY8 (Chen et al., 2010; Ren et al., 2024). Since WRKY48 and WRKY8 belongs to the subgroup class IIc, we also chose WRKY28, another member of class IIc, to check their binding efficiency with *CIPK6* promoter. *CIPK6* upstream region has two putative WRKY binding sites, TTGAC at -228 bps (P1) and GGTCA at -268 bps (P2) upstream from the transcription start site (Figure 1A). Initially, we examined the binding of the three WRKY proteins (WRKY48, WRKY28, and WRKY8) to the *CIPK6* promoter. We used oligonucleotides (∼100bps) containing both the binding sites (P1 and P2) and purified the WRKY-HIS fusion proteins in electromobility shift assay. The only WRKY protein that exhibited binding activity on the *CIPK6* promoter was WRKY48 (Figure 1B, Supplementary Figure 1A). WRKY48 binding to specific cis-acting element was verified using probes with substituted bases. We further looked for its differential binding activity to the P1 or P2 binding element. For P1 and P2, we independently designed oligonucleotides about 40 bps long, and examined the binding activity of WRKY48 (Figure 1C) as well as WRKY28 and WRKY8 (Supplementary Figure 1A). We observed that the WRKY48 binding activity was higher on the P1 binding element than on the P2 binding element. For WRKY28 and WRKY8 we could not see the binding activity on either P1 or P2 element (Supplementary Figure 1A). The results suggest, WRKY48 specifically binds to the *CIPK6* promoter.

**Figure 1.**
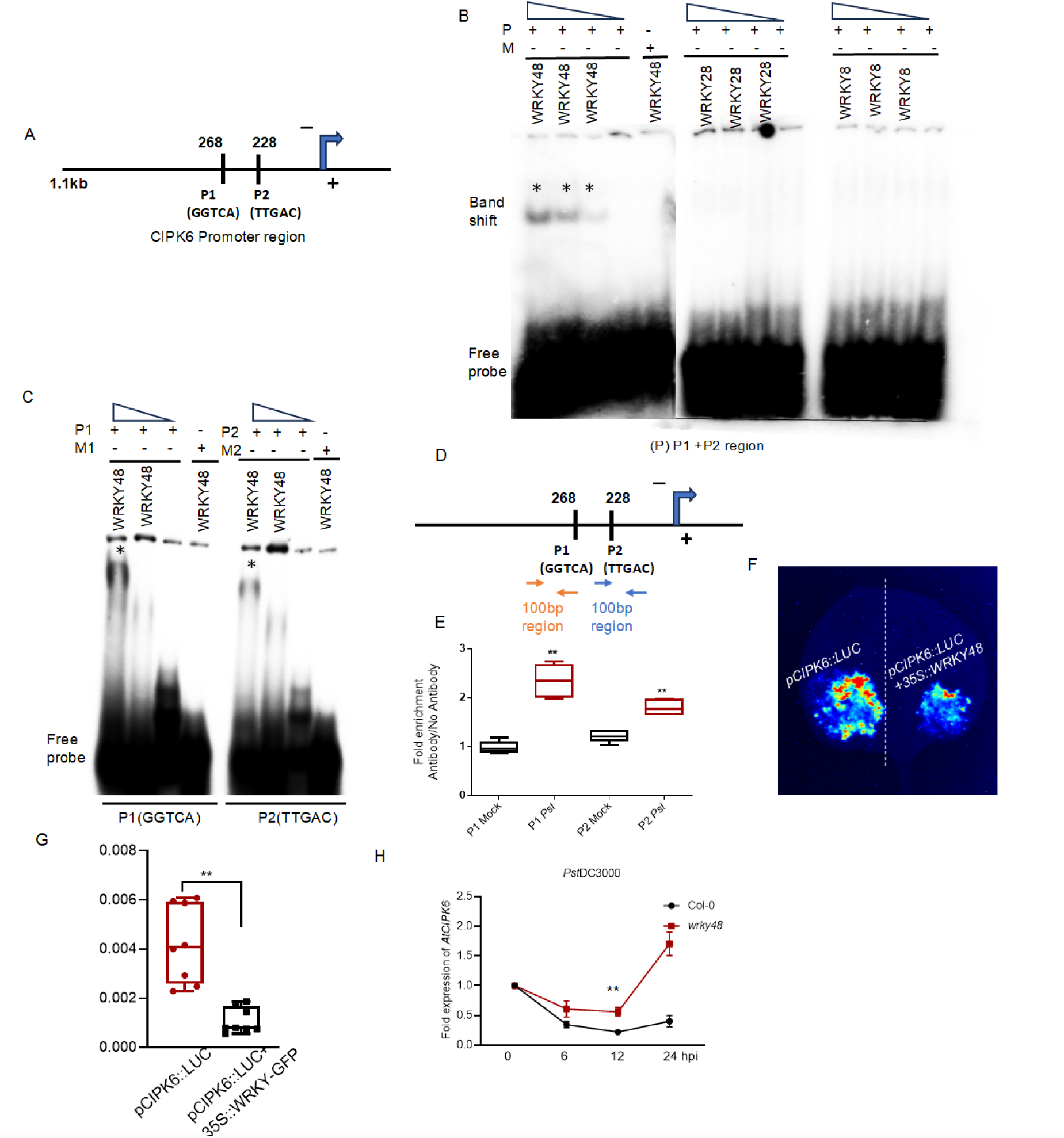
WRKY48 binds to *CIPK6* promoter and it negatively regulate *CIPK6* expression during *Pst*DC3000 challenge (A) Schematic representation of WRKYs (48, 28, and 8) transcription factor binding sites on *CIPK6* promoter. (B) EMSA to show the binding of WRKY 48, WRKY28, and WRKY8 on the *CIPK6* promoter using 4µg, 2µg and 1µg protein concentrations of each. (C) EMSA showing that WRKY 48 has greater binding on P2 region of the CIPK6 promoter using 2ug of WRKY48 protein concentration whereas probe was prepared by using 100ng/ul for each oligo ( sense or antisense). (D) schematic diagram of the region on the *CIPK6* promoter for pCHIP-qPCR (E) Fold Enrichment of WRKY48 (GFP-Ab) on P1 and P2 region of *CIPK6* promoter during *Pst*DC3000 challenge. Data are ±SD of 2 biological samples with each of them has three technical replicates, P>0.001 (F) Qualitative luciferase reporter assay in *Nicotiana benthamiana* leaves. Luciferase assay has been performed and image has been captured after 60 hours of Agrobacterium mediated co-infiltration of *pCIPK6:LUC* construct and *pCIPK6::LUC* +*35::SWRKY48* constructs. (G) dual luciferase assay has been performed in *Nicotiana* leaves to investigate the transcriptional regulation of CIPK6 in the presence of WRKY48. Data points on the bar shows each biological replicates and data are the means of ±SD of 8 biological samples,P>0.005. (H) Expression of *CIPK6* in Col-0 and *wrky48* mutant at the indicated time points (0h, 6h,12h, 1h and 24h post inoculation of *Pst*DC3000. Data are means of ±SD of 3 biological samples with each of them has two technical replicates, with significance of **P≥0.001 and *P ≥ 0.05.

### WRKY48 negatively regulate transcription of *CIPK6* during *Pst*DC3000 infection

To elucidate the possible effect of the association of WRKY48 with *CIPK6* promoter during pathogenesis we compared the relative occupancy of WRKY48 protein on *CIPK6* promoter between mock and *Pst*DC3000-challenged WRKY48-GFP overexpressing plants by ChIP--qPCR following a 12-hour *Pst*DC3000 inoculation. When compared to mock-treated plants, we found that *Pst* inoculation significantly increased WRKY48-GFP binding in the *CIPK6* promoter at both P1 and P2 binding region. As expected, the relative abundance of WRKY48-GFP was more at P1 region as compared to P2 during *Pst* challenge (Figure 1D and E). The equivalent expression of *WRKY48-GFP* in the *35S::WRKY48-GFP* plants were confirmed through western blot (Supplementary figure 1B). We performed dual luciferase assay to study the transcriptional regulation of CIPK6 by WRKY48. We utilised a promoter:reporter construct by putting the *CIPK6*-promoter (∼1.2Kb) upstream to the firefly luciferase gene *LUC* in the dual luciferase reporter assay by transiently expressing effector (35S::WRKY48-GFP) and reporter together in *Nicotiana benthamiana* leaves. We observed that WRKY48 repressed Luciferase activity (Figure 1F). A quantitative assay of relative luciferase activity in terms of luminescence produced with respect to constitutively expressing Renilla-LUC in presence of WRKY48 showed a 3-fold reduction as compared to the sample lacking *35S::WRKY48-GFP* (Figure 1G).

To corroborate this result, we examined the *CIPK6* transcript accumulation in Col-0 and *wrky48* mutant plants at the denoted time points following *Pst*DC3000 inoculation at 5x10^5^ CFU/ml. *Pst* inoculated Col-0 plants displayed higher CIPK6 transcript accumulation as compared to *wrky48* mutant (Figure 1H). The results altogether suggest WRKY48 negatively regulate CIPK6 expression during bacterial pathogenesis in Arabidopsis.

### WRKY48 negatively impact plant defense associated with reduced ROS generation in pattern triggered immunity

To get insight into the role of WRKY48 in plant defense, we first examined the transcript profile of *WRKY48* after *Pst*DC3000 challenge. qPCR data showed enhanced accumulation of WRKY48 mRNA in Col-0 plants at the early hours, however, the peak declines after 90 min of *Pst* challenge (Figure 2A). Reduced bacterial growth was observed in the *wrky48* mutant as compared to that in Col-0, although both the lines showed significantly increased bacterial growth with respect to *cipk6* mutants (Figure 2B).

**Figure 2.**
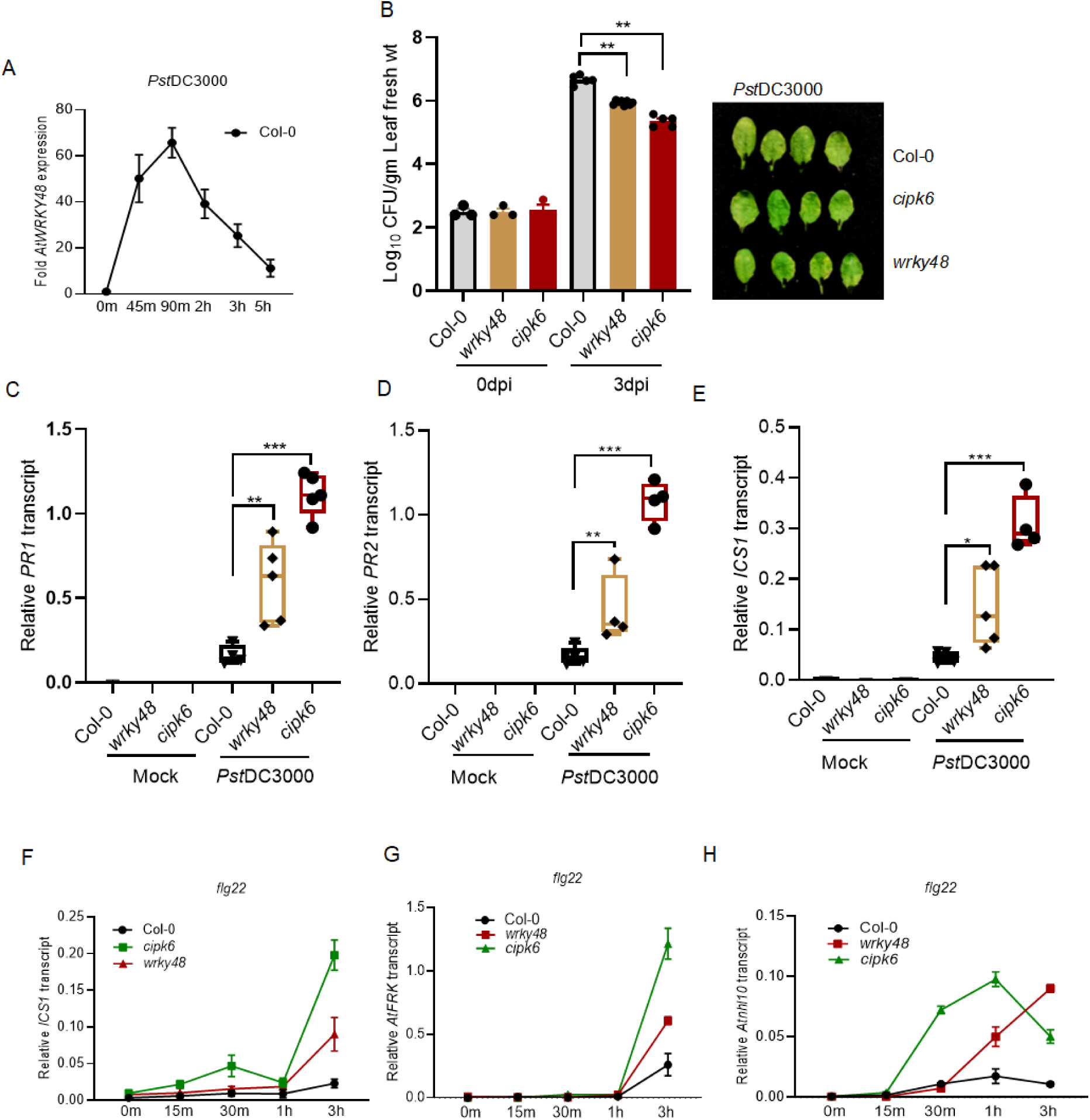
WRKY48 negatively impact pattern triggered immunity. **(A)** WRKY48 expression is induced at the initial indicated time points during *Pst*DC3000 challenge in 5-week-old leaves of Col-0. **(B)** Bacterial Load and disease symptoms in Col-0, *wrky48* mutants, and *cipk6* mutants at 0 dpi and 3 dpi. **(C)**, **(D)**, and **(E)** transcript accumulation of *PR1*, *PR2*, and *ICS1* respectively in mock and *Pst*DC3000 challenged leaves tissues of Col-0, *wrky48*, and *cipk6* mutants at 12hpi. **(F)**, **(G)**, and **(H)** transcript accumulation of *ICS1* and PTI markers *AtFRK* and *Atnhl10* respectively in 5-week-old leaves of Col-0, *wrky48*,and *cipk6* mutants at the denoted time points after flg22 application. Data are mean of ± SD of three biological samples, each biological replicate has two technical replicates with with **P≥0.001 and *P ≥ 0.05. Experiments are repeated twice with similar results.

In agreement with this observation and negative role of WRKY48 in plant defense, *wrky48* mutants showed higher expression of defense marker genes like *PR1*, *PR2*, and *ICS1* as compare to Col-0 after *Pst* infiltration, whereas in comparison to *cipk6* mutants, the expressions were significantly lower at the indicated time points (Figure 2C, D, and E). Further, to investigate the role of WRKY48 in PTI responses, 5µM flg22, a PTI elicitor, was infiltrated into the Col-0, *cipk6* and *wrky48* mutant leaves to monitor activation of PTI response by assessing transcript accumulation of the PTI markers e.g. *ICS1*, *nHL10*, and *FRK1*. The expression of *ICS1*, *nHL10*, and *FRK1* was considerably greater in *cipk6* mutants than in Col-0 and *wrky48* mutants (Figure 2F, G, and H).

The production of reactive oxygen species (ROS) is essential for successful activation of plant immunity (Lee et al., 2020; Dumanović et al., 2021). In order to ascertain the involvement of WRKY48 in ROS generation during PTI activation, Col-0, *wrky48*, and *cipk6* mutants were challenged with *Pst*DC3000, followed by DAB staining and a quantitative Luminol-based assay. The *wrky48* mutant displayed significantly less ROS generation (Figure 3A, Supplementary Figure 4A), less DAB staining (Figure 4B and Supplementary Figure 4B). The expression of ROS scavenging genes such as, *CAT1* and *APX1* was higher in the *wrky48* mutant than in the *cipk6* mutant. Significantly more intense DAB staining, more ROS generation, and less expression of *CAT1* and *APX1* were exhibited by *wrky48* than the Col-0 (Figure 3C and D) suggesting WRKY48 and CIPK6 both are essential for normal oxidative burst during PTI.

**Figure 3.**
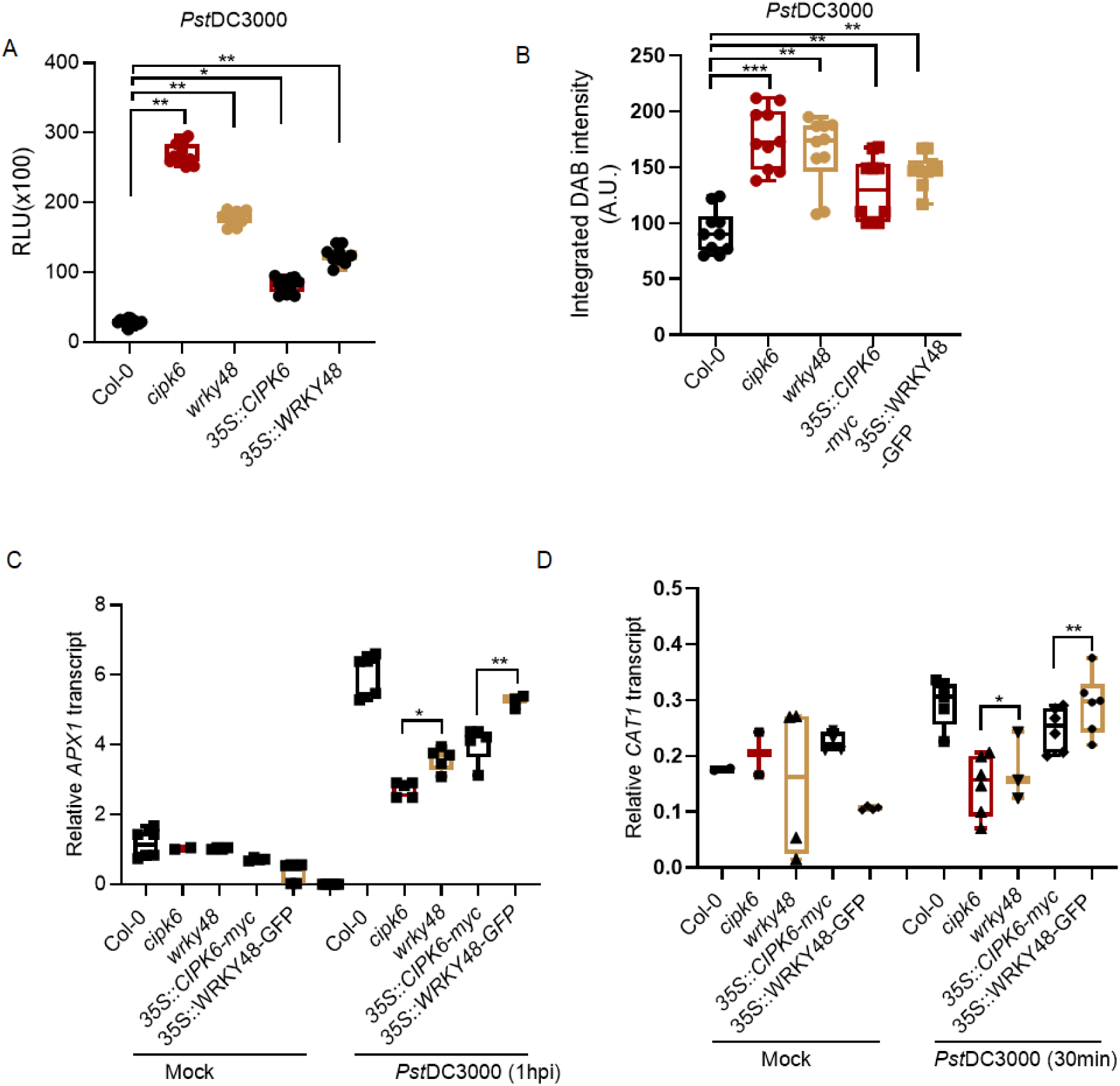
wrky48 mutants show enhanced ROS generation as compare to Col-0; however, when compared to cipk6 mutant, ROS is decreased. **(A)** Box-plot displaying the highest ROS produced (∼10-12 m) during the course of 60 m after PstDC3000 inoculation and starting the quantification based on the luminescence produced after the reaction of H2O2 and Luminol in the presence of Horse Radish Peroxidase (n=12 leaf discs for each genotype and data are ± SD of mean values, *P≥0.05 or **P ≥ 0.001). **(B)** DAB intensity 4hpi in Col-0, cipk6, wrky48, 35S::CIPK6-myc, and 35S::WRKY48-GFP. Number of leaves used for the analysis of DAB intensity 8, data plotted is ± SD of mean associated values for each genotype. **(C)** and **(D)** Transcript accumulation of ROS markers, APX1 and CAT1 in Col-0, cipk6, wrky48, 35S::CIPK6-myc, and 35S::WRKY48-GFP in mock and PstDC3000 inoculated leaves at indicated time points. Data are ±SD of 3 biological samples with each of them has three technical replicates, with **P≥0.001 and *P ≥ 0.05.

**Figure 4.** WRKY48 interacts with MPK3 during PstDC3000 challenge. **(A)** Yeast two hybrid assay to show the interaction between MPK3 and WRKY48. pGBKT7-WRKY48 and pGADT7-MPK3 was co-transformed in Y2H gold yeast cells and colonies were further serially diluted and allowed to grow on selective synthetic medium deficient in Leu, Trp, His, and Ade with x-ɑgal to check the GAL4 reporter activity. **(B)** BIFc assay to check the interaction after transient expression of cd31648-WRKY48 and cd31651-MPK3, co-infiltrated in *Nicotiana* leaves. The reconstitution of YFP was seen at 48 post hours infiltration. **(C)** GST Pull down assay between MPK3-GST and WRKY48-6xHis to check interaction *in vitro*. **(D)** Split luciferase assay to check the interaction in *vivo* between MPK3-nluc-pCAMBIA1300 and WRKY48-cluc-pCAMBIA1300 in *Nicotiana* leaves. **(E)** Co-Immunoprecipitation in *35s::WRKY48-GFP* Arabidopsis plants to check the interaction between MPK3 and WRKY48, immunoprecipitation was done by MPK3-Ab followed by immunoblotting with MPK3-Ab and GFP-Ab. **(F)** Co-Immunoprecipitation was also done in Arabidopsis in *Pst*DC3000 challenged *35S::WRKY48-GFP* and *35S::WRKY48-GFP/mpk3* plants.

### WRKY48 interacts with MPK3 during pathogenesis

The induction of PTI is widely recognized to involve the concurrent activation of MAP kinases (MPKs) and Ca²⁺-dependent protein kinases (CPKs). Prior research has demonstrated that pathogen-induced activation of CPK5/7/11 leads to the phosphorylation of WRKY48, WRKY28, and WRKY28 which subsequently mediate downstream transcriptional reprogramming (Gao et al., 2013). In order to define the role of MPK3/6-mediated phosphorylation of WRKY48 in the negative regulation of PTI, we sought to understand the relationship between MPK3/6 and WRKY48. For this, first we checked the direct interaction between MPK3/6 with WRKY48 through yeast two hybrid assay. Both MPK3 (Figure 4A) and MPK6 (Supplementary Figure 2A) exhibited the interaction with WRKY48 in yeast. We then examined the direct interaction between the proteins by in vitro GST pull down assay by incubating purified recombinant GST-MPK3 or GST-MPK6 proteins with His-WRKY48 protein. Interaction between MPK3 and WRKY48 was visible in the western blot (Figure 4C) while, the interaction between MPK6 and WRKY48 could not be detected in this assay (Supplementary Figure 2B)).

We further validated *in planta* interaction between MPK3 and WRKY48 using BIFc assay and split luciferase assay in *Nicotiana benthamiana* and co-immunoprecipitation assay using Arabidopsis overexpressing lines. In BiFC assay we could visualize the interaction between MPK3 and WRKY48 most likely in the nucleus (Figure 5B). while the co-expression of cd3-1648-MPK6 and cd3-a651-WRKY48 did not show YFP fluorescence (Supplementary Figure 2E). In the split-luciferase experiment also, the transient expression of pCAMBIA1300-nluc-MPK3 and pCAMBIA1300-cluc-WRKY48 in *Nicotiana benathamiana* leaves exhibited much higher intensity of relative chemiluminescence as compared to the sample co-expressing pCAMBIA1300-nLUC and WRKY48 (Figure 4D). This suggests that MPK3 interacts with WRKY48 *in vivo.* To further ascertain this interaction in Arabidopsis, we did the co-immunoprecipitation assay using the cell extract of the Arabidopsis plant co-expressing *35S::WRKY48-GFP/wrky48* and *35S::WRKY48-GFP/mpk3*. MPK3 protein was immunoprecipitated with MPK3 specific antibody followed by immunoblot with GFP antibody and MPK3 antibody. We could detect the interaction between MPK3 and WRKY48 in the plant protein expressing both WRKY48-GFP and MPK3, whereas we could not detect the presence of GFP-specific band where WRKY48-GFP was expressed under *mpk3* mutant background, suggesting MPK3 and WRKY48 interact in Arabidopsis (Figure 4E).

**Figure 5.**
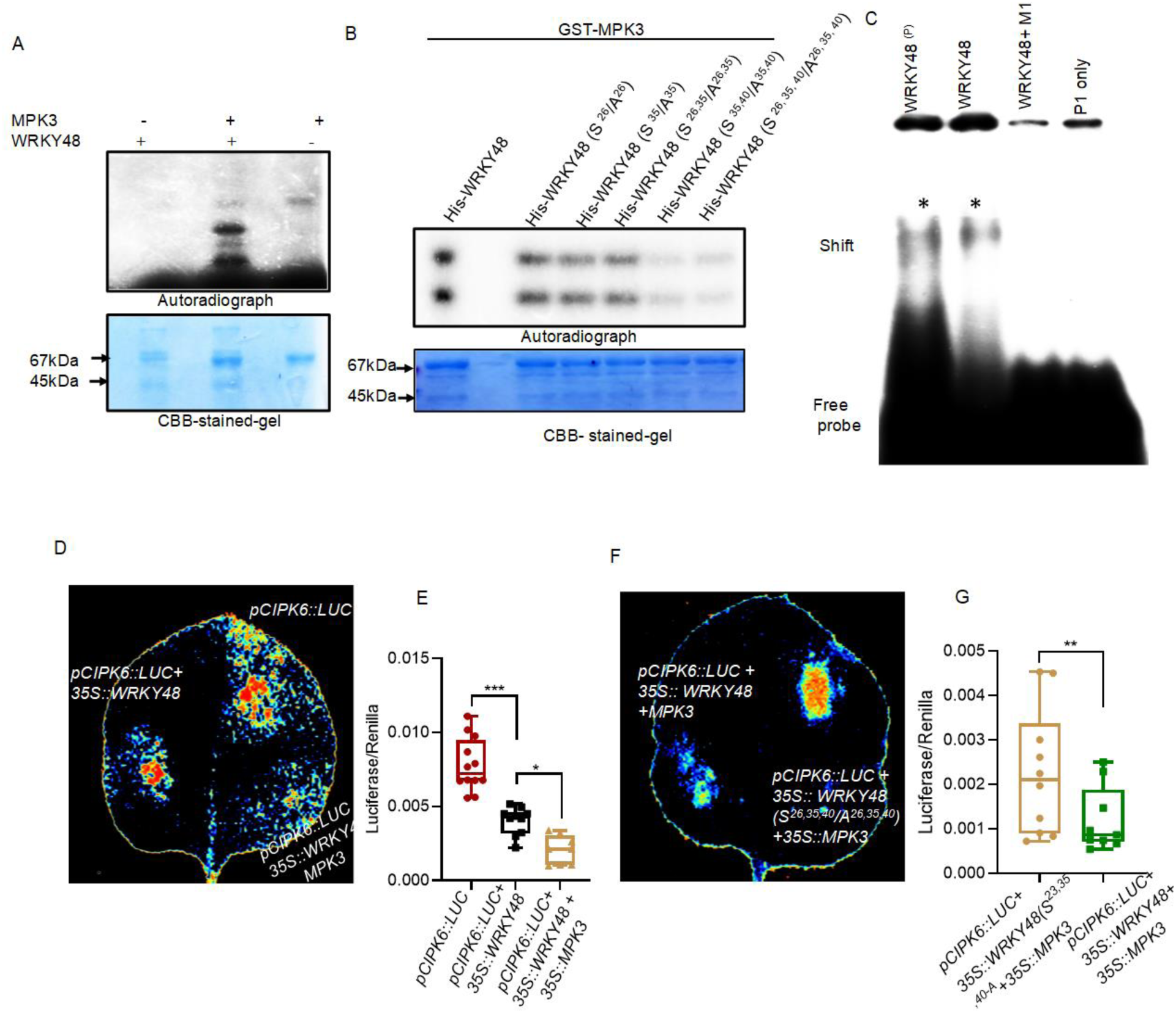
WRKY48 binding on *CIPK6* promoter is dependent on its phosphorylation through MPK3. **(A)** *in vitro* phosphorylation of WRKY through MPK3. **(B)** *in vitro* phosphorylation of WRKY48 Serine (S^26^, S^35^, and S^40^) mutants through MPK3. **(C)** EMSA to check the binding of phosphorylated WRKY48 on *CIPK6* promoter. **(D)** and **(E)** qualitative luciferase assay in *Nicotiana benthamiana* leaves. Luciferase assay has been performed and image has been captured after 60 hours of Agrobacterium mediated co-infiltration of *pCIPK6:LUC + 35S::WRKY48-GFP and pCIPK6::LUC+35::SWRKY48+ 35S::MPK3-myc* constructs in one leaf and in another leaf *pCIPK6::LUC+35::SWRKY48+ 35S::MPK3-myc and pCIPK6::LUC+35S::WRKY48(S^26,35,40^/A)+ 35S::MPK3-myc* **(F)** and **(G)** dual luciferase assay has been performed in *Nicotiana* leaves to asses the transcriptional regulation of CIPK6 in the presence of MPK3 and WRKY48, and *WRKY48(S^26,35,40^/A)+*. Data points on the bar shows each biological replicates and data are the means of ±SD of 8 biological sample, P>0.005.

To investigate the biological significance of MPK3 and WRKY48 interaction in Arabidopsis during a *Pst*DC3000 challenge, we treated plants overexpressing WRKY48 in the backgrounds of *wrky48* mutant or *mpk3* mutant with 10 mM MgCl₂ as a mock control or inoculated them with *Pst*DC3000. Proteins were extracted 3 hours post-inoculation, immunoprecipitated using MPK3-specific antibody followed by immuno blot with MPK3 and GFP antibodies (Figure 5F). The results showed a significantly enhanced interaction between MPK3 and WRKY48 in the *Pst*DC3000-treated plant as compared to the mock samples. These findings indicate that the bacterial infection increases MPK3 and WRKY48.

### Phosphorylation of WRKY48 by MPK3 enhances its binding affinity to the CIPK6 promoter and thereby promotes trans-repressive activity

In our previous results, we demonstrated the interaction between MPK3 and WRKY48. To examine the possibility of MPK3-mediated phosphorylation of WRKY48, we incubated recombinant WRKY48-His protein with MPK3-GST or MPK3kd-GST (kinase dead) in a kinase assays. Additionally, we conducted a kinase assay using MPK6-GST and WRKY48-His. As expected, WRKY48 exhibited phosphorylation only in presence of MPK3-GST (Figure 6A), while MPK6-GST showed only autophosphorylation (Supplementary Figure 2C). MPKs are serine/threonine-specific protein kinases, which preferably phosphorylate the residues at the C-terminus of proline (Pearson et al., 2001; Meng and Zhang, 2013b). WRKY48 contains three such serine residues (S26, S35, and S40) and one threonine residue (T16), which are conserved among the dicots such as *Arabidopsis*, *Brassica*, *Raphanus*, and *Camelina*, as well as in the monocot maize (Supplementary Figure 3).

**Figure 6.**
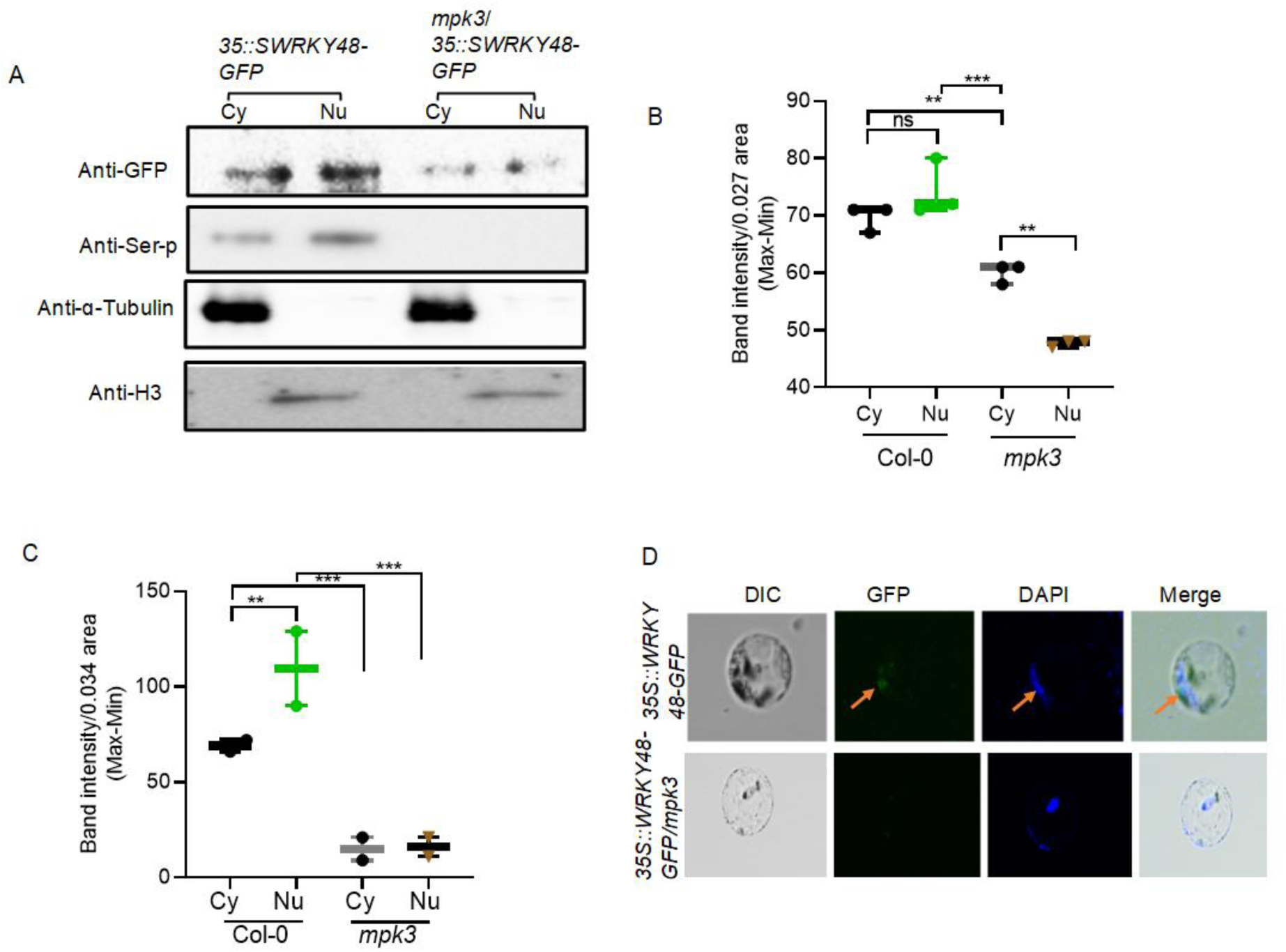
Nuclear import of WRKY48 is associated with its increased phosphorylation. **(A)** Phosphorylation of WRKY48 through MPK3 is associated with its nucleo-cytoplasmic distribution. The WRKY48-GFP fusion was overexpressed in Col-0 and mpk3 mutant as described above and nuclear and cytoplasmic protein fraction extracted from these plants were immunoprecipitated with GFP-Ab and detected in immunoblot developed through GFP–Ab in cytoplasmic and nuclear fraction. To check the phosphorylation of WRKY48 the same immunoblot was also developed through Ser-P-Ab, ɑ-Tubulin-Ab for cytoplasmic fraction, and Histone3 for nuclear fraction. **(B)** and **(C)** to quantify the band intensity of WRKY48-GFP band intensity in immunoblot developed through GFP-Ab in nuclear and cytoplasmic fraction and phosphorylation of WRKY48-GFP respectively. The experiment was repeated thrice for nucleo-cytoplasmic distribution of WRKY48 and twice to check the phosphorylation. The data for band intensity is ±SD of mean of three and two biological samples for each fraction, where **P≥0.001 and ***P≥0.0001, for significance **(D)**. Nuclear localisation of WRKY48.The WRKY48-GFP fusion was overexpressed driven by the *CaMV 35S* promoter in Col-0 and *mpk3* mutant plants The brown arrow indicate the expression of WRKY48 in the nucleus, DAPI used for nucleus staining.

To investigate the serine phosphorylation activity of MPK3, we generated phospho-null mutants of WRKY48 by replacing serine residues with alanine, either individually or in combinations (S^26^/A^26^, S^35^/A^35^, S^26^S^35^/A^26^A^35^, S^35^S^40^/A^35^A^40^, and S^26^S^35^S^40^/A^26^A^35^A^40^). These recombinant mutated proteins were used as substrates for MPK3-GST as kinase in the kinase reactions. Individual substitution of S^35^ significantly reduced WRKY48 phosphorylation, whereas substitution of S^26^ had no substantial effect. However, combined replacement of S^35^S^40^ or S^26^S^35^S^40^ with alanine drastically reduced phosphorylation, suggesting that S^35^ and S^40^ are the primary MPK3-mediated phosphorylation sites (Figure 6B).

To examine the impact of MPK3-mediated phosphorylation of WRKY48 on its binding affinity to the *CIPK6* promoter, the MPK3-phosphorylated and unphosphorylated WRKY48 proteins were used in a EMSA assay with a radiolabelled *CIPK6* promoter probe containing the P1 and P2 regions. Interestingly, phosphorylated WRKY48 exhibited significantly higher binding affinity compared to its unphosphorylated form (Figure 6C).

To determine the impact of MPK3-mediated phosphorylation of WRKY48 on its trans-repressive activity towards the CIPK6 promoter, we conducted a dual luciferase reporter assay by transiently co-expressing effector and reporter constructs in *Nicotiana benthamiana* leaves. The reporter and effector constructs were tested in three combinations: *pCIPK6::LUC* only, *pCIPK6::LUC* + *35S::WRKY48-GFP*, and *pCIPK6::LUC* + *35S::WRKY48-GFP* + *35S::MPK3-GFP*. Luciferase activity, measured as chemiluminescence, was observed two days post-infiltration. The results indicated that WRKY48, in the presence of MPK3, significantly repressed luciferase expression as compared to its absence (Figure 6D). To confirm these findings, we performed a quantitative dual luciferase assay by normalizing relative luciferase activity (luminescence) to the constitutive expression of Renilla-LUC. The relative luminescence was significantly lower (Figure 6E) in the presence of WRKY48 and even more so when MPK3 was co-expressed, as compared to the control having only the reporter construct (*pCIPK6::LUC*).

Further, when WRKY48 was substituted at all the three serine residues (S^26^ ^35^ ^40^/A^26^ ^35^ ^40^) with alanine, the repression of *pCIPK6::LUC* activity by WRKY48 was abolished, restoring luciferase expression levels at the level of the control (Figure 6F). This suggests that WRKY48 after phosphorylation by MPK3 exhibits higher binding affinity to the *CIPK6* promoter and effectively represses its activity, whereas unphosphorylated WRKY48 binds to CIPK6 promoter with lower affinity and does not repress its activity as efficiently (Figure 6G). Altogether, results suggest, MPK3-mediated phosphorylation of WRKY48 enhances its binding affinity to the *CIPK6* promoter, thereby strengthening its trans-repressive activity.

### The phosphorylated WRKY48 is predominantly present in the nucleus

Post-translational modification of proteins influences their structure, localization, and numerous cellular processes (Friso and Van Wijk, 2015). In our study, we observed that phosphorylated WRKY48 binds to the *CIPK6* promoter with increased affinity. We aimed to determine whether MPK3-mediated phosphorylation is necessary for the nuclear localization of WRKY48 since, BiFC results showed that MPK3 preferentially interacts with WRKY48 in the nucleus. To investigate this, we extracted the cytoplasmic and nuclear fractions from *WRKY48-GFP*/*wrky48* and *WRKY48-GFP/mpk3* plants and performed immunoblotting using an anti-GFP antibody. Our results revealed that significantly less WRKY48 protein was detected in the nuclear fraction in absence of MPK3 as compared to the nuclear fraction of *35S::WRKY48-GFP*/*wrky48* plants (Figure 7A, B). To confirm whether the nuclear-localized WRKY48 is phosphorylated, we immunoblotted the same subcellular fractions from the previous experiment using a phospho-serine antibody. We found that both cytoplasmic and nuclear WRKY48 were phosphorylated, but the nuclear-localized WRKY48 exhibited greater phosphorylation than the cytoplasmic fraction (Figure 7A, C). The MPK3-mediated phosphorylation of WRKY48 and its nuclear localization were further confirmed by the nuclear localization of WRKY48 when the WRKY48-GFP fusion protein was overexpressed under the control of the *CaMV35S* promoter in both Col-0 and mpk3 mutant plants (Figure 7D). The observation that MPK3-mediated phosphorylation of WRKY48 is predominantly localized in the nucleus indicates that this post-translational modification plays a crucial role in regulating function of WRKY48 within the nucleus.

**Figure 7.**
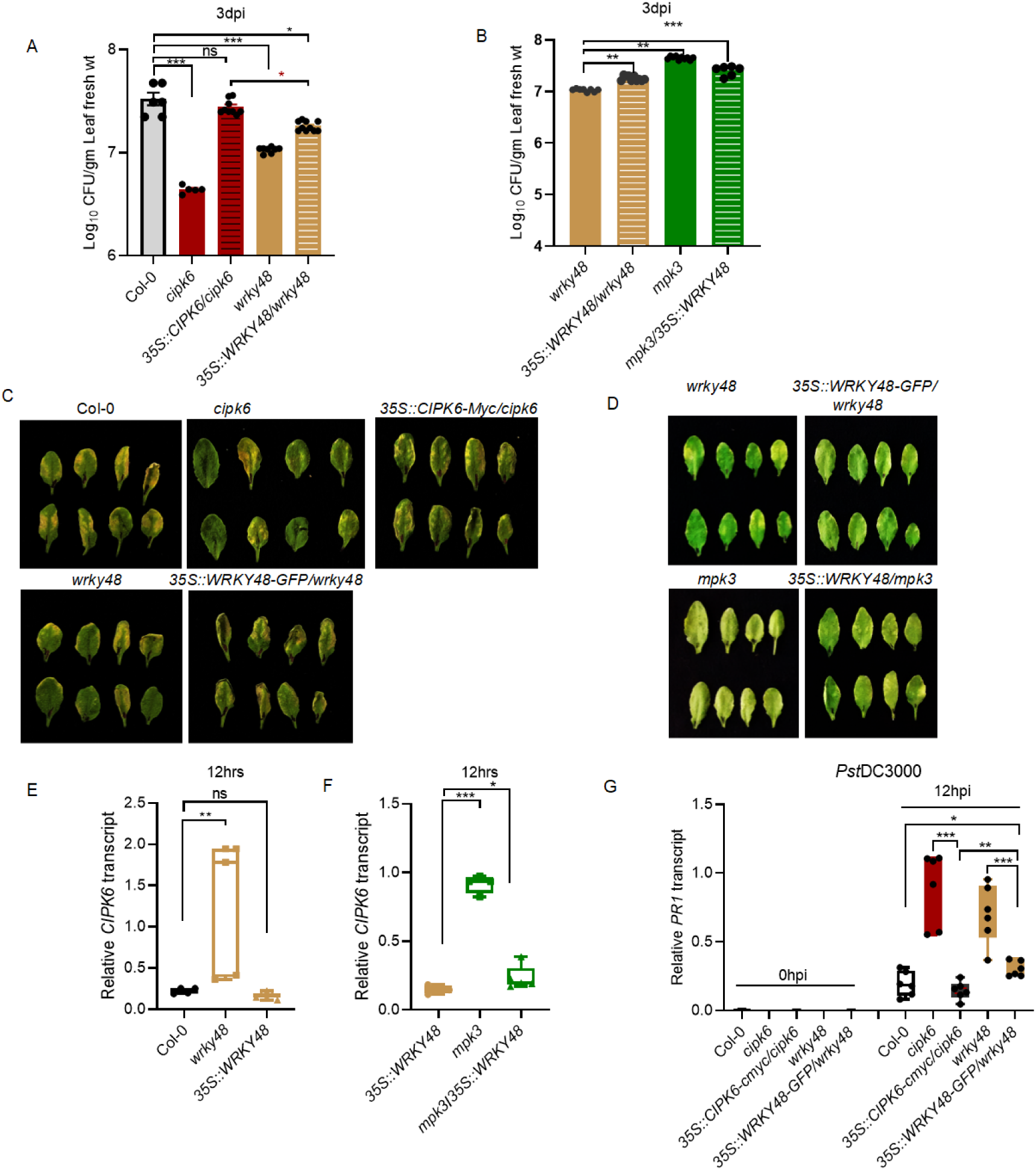
Constitutive expression of WRKY48 results in reduced transcript accumulation of *CIPK6*. (A) Bacterial count in Col-0, *cipk6*, *wrky48, 35S::WRKY48,* (B) Bacterial load in *wrky48, 35S::WRKY48-GFP/wrky48, mpk3, and 35S::WRKY48-GFP/mpk3,* the bacterial count data are statistically significant with ± SD of 8 biological replicates with ***P ≥0.0001, **P≥0.001 and *P ≥ 0.05. CFU count data for *wrky48* mutant and *35S::WRKY48-GFP* is common for both figure A and B, since the single experiment was performed and the data has been split to gain visibility of the significance. (C) *and* (D) *Pst* induced symptoms of indicated genotypes at 3dpi (E) Relative CIPK6 transcript in Col-0, *wrky48, and 35S::WRKY48* (F) Relative *CIPK6* transcript in *mpk3* and *35S::WRKY48.* In figure E and F data for 35S::WRKY48 is common, since it was a single experiment (G) Relative *PR1* transcript in Col-0, *cipk6*, *35S::CIPK6-myc*, *wrky48*, *35S::WRKY48-GFP.* qPCR data are ±SD of 3 biological samples with each of them has three technical replicates, with ***P ≥0.0001, **P≥0.001 and *P ≥ 0.05.

### WRKY48 phosphorylation is induced by MPK3 in response to the *Pst*DC3000 challenge, leading to its enhanced trans-repression of the *CIPK6* transcript

Phosphorylation of a protein by Mitogen-Activated Protein Kinase (MAPK) can result in a protein’s degradation, stabilization, interaction with other proteins or translocation, depending on the specific protein and the phosphorylation site that in turn promote disease resistance through activation of downstream genes (Plotnikov et al., 2011; Zhang and Zhang, 2022).The protein phosphorylation is essential phenomena for plant defence, since it controls the signalling pathways that trigger the immunological response against pathogens (Park et al., 2012a). For instance, Arabidopsis CAMTA3, a TF known to be a negative regulator of plant defense, is phosphorylated by MPK3 and MPK6 after flg22 application. The phosphorylation of CAMTA3 results in its destabilisation and promote its nuclear import (Jiang et al., 2020). In our study, we investigated the role of MPK3-mediated phosphorylation of WRKY48 in plant immunity. To this end, we first overexpressed WRKY48 in *wrky48* mutant and *mpk3* mutant background, the homozygous lines were screened and confirmed in T2 generation through semi-qPCR and immunoblot analysis (Supplementary Figure 4A and B). We examined the defense responses of Col-0, *35S::WRKY48-GFP*/*wrky48*, and *35S::CIPK6-myc*/*cipk6* plants following a challenge with *Pseudomonas syringae* pv. *tomato* DC3000 (*Pst*DC3000). After 3 days post-inoculation (dpi), bacterial loads were significantly higher in *35S::CIPK6-myc*/*cipk6* plants compared to *35S::WRKY48-GFP*/*wrky48* plants (Figure 7A). Overexpression of *WRKY48* in the *mpk3* mutant background rendered the plants even more susceptible than *35S::WRKY48-GFP*/*wrky48* plants (Figure 7B). Interestingly, overexpression of *WRKY48* resulted in significantly lower expression of *CIPK6* compared to *wrky48* mutants, with levels similar to Col-0 plants at 12 hours post *Pst*DC3000 challenge (Figure 7C). In contrast, *CIPK6* expression in *mpk3* mutants was significantly higher than in *WRKY48*-overexpressing plants (Figure 7D). These findings suggest that both MPK3 and WRKY48 are required to suppress *CIPK6* transcript levels, thereby enhancing resistance to bacterial growth. Consistent with these results, the expression of *PR1* was significantly higher in *35S::WRKY48-GFP* plants compared to *wrky48* mutant and in *35S::CIPK6-myc* plants compared to *cipk6* mutants (Figure 7E).

Now, to determine whether MPK3-mediated phosphorylation of WRKY48 is triggered by *PstDC3000* challenge in planta, we analyzed the in vivo phosphorylation of WRKY48 in *35S::WRKY48*/wrky48 and *35S::WRKY48*/*mpk3* plants 60 minutes post-challenge. Our results showed that WRKY48 phosphorylation was induced following the pathogen challenge in *35S::WRKY48*/*wrky48* plants (Figure 8B). However, phosphorylation levels were significantly lower and remained unchanged after the pathogen challenge in *35S::WRKY48*/*mpk3* plants. Additionally, during a *Pst* challenge, the expression of MPK3 increases up to 1 hpi, and in conjunction with this, the accumulation of WRKY48 is maintained, showing enhancement until 3 hours post-inoculation. It indicates *Pst* challenge triggers the MPK3 mediated phosphorylation of WRKY48 in *planta* (Figure 8A). Further, to investigate whether WRKY48 also regulates CIPK6 at the translational level, we expressed *pCIPK6::CIPK6-myc* under *35S::WRKY48*/*wrky48*, *wrky48*, and *35S::WRKY48*/*mpk3* backgrounds. CIPK6 protein levels were then analyzed following a *Pst* DC3000 challenge at specified time points using myc-antibody. We observed that the CIPK6 protein level decreased in *35S::WRKY48*/*wrky48* plants following a *Pst*DC3000 challenge, whereas it was significantly elevated in the *wrky48* mutant 45 minutes post-inoculation (Figure 8C). In *35S::WRKY48*/*mpk3* plants, increase in CIPK6 protein level was delayed until 1 hours post-inoculation (Figure 8C). Additionally, overexpression of *CIPK6* in the *wrky48* mutant background led to a significant reduction in bacterial load compared to plants constitutively expressing *CIPK6* (Figure 8D). This was further supported by the disease symptoms observed following *Pst* infection (Figure 8E). The above findings collectively imply that MPK3 is required for the phosphorylation of WRKY48, that in turn negatively regulates CIPK6 transcription and translation, contributing to plant resistance.

**Figure 8.**
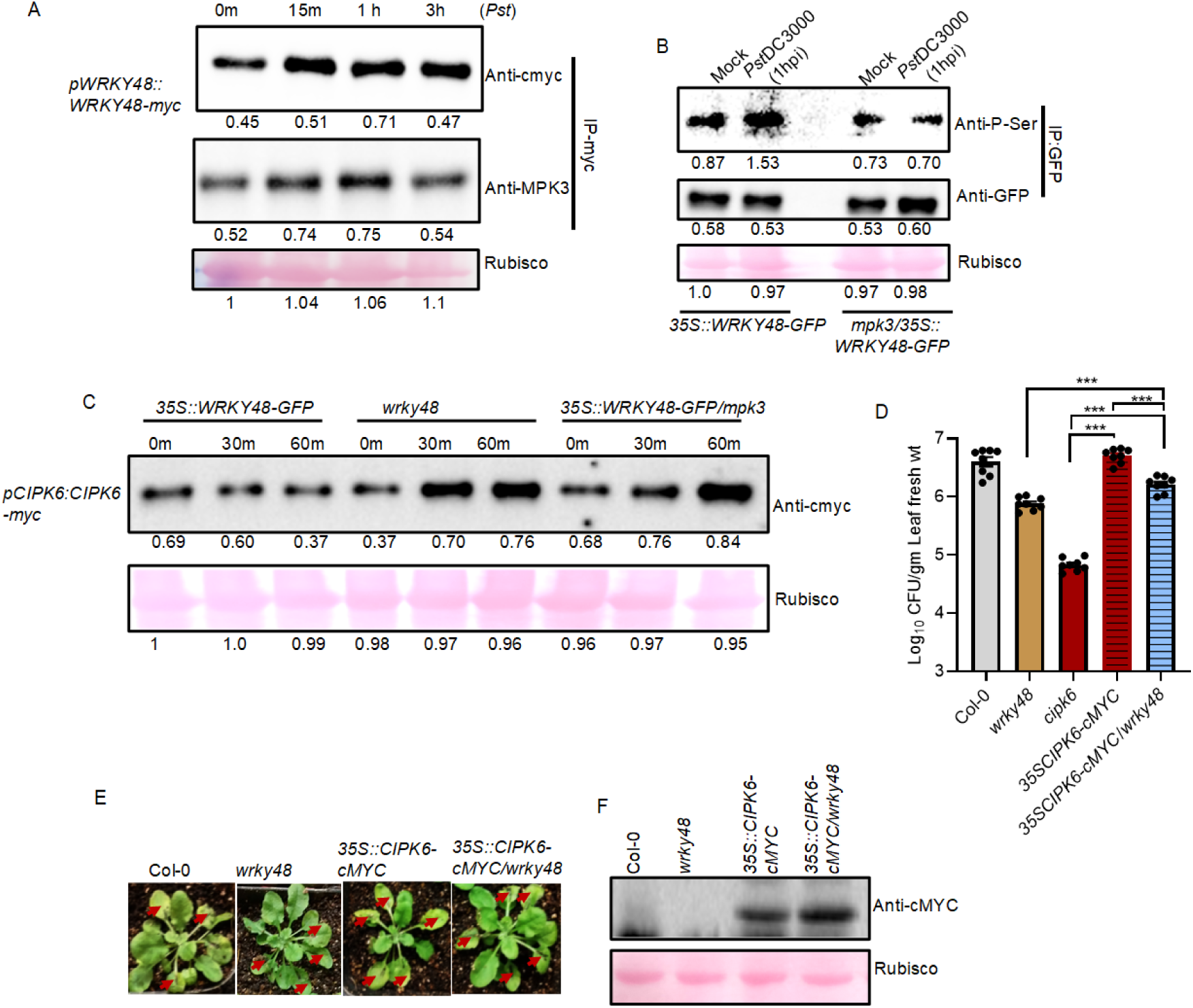
Expression of WRKY48 is induced during *Pst*DC3000 challenge and it is phosphorylated through MPK3 in planta that in turn regulate the translation of CIPK6. **(A)** *pWRKY48::WRKY48-myc* expressing plants were infiltrated with *Pst*DC3000 was and leaves tissue was harvested at the denoted time points, equal amount of crude protein was immunoprecipitated with cmyc-Ab and western blot was developed through myc, MPK3, and GST antibody. Band intensity is denoted with respect to loading control Rubisco, below the blot **(B)** *in planta* phosphorylation of WRKY48 (band intensity with respect to loading control) during pathogen challenge through MPK3 was done in *35S::WRKY48-GFP* and *35S::WRKY48-GFP/mpk3* expressing stable Arabidopsis plants. Equal concentration of total protein was used to do the western through Ser-Ab and GFP-Ab, ponceauS stain for equal protein loading. Band intensity (denoted by number) of phosphorylation with resect to WRKY-GFP band intensity/protein concentration **(C)** *pCIPK6::CIPK6-myc* expressed in *35S::WRKY48-GFP*, *wrky48*, and *35S::WRKY48-GFP/mpk3* background, After *Pst*DC3000 infiltration leaves tissue was harvested at the indicated time points, equal amount of crude protein was immunoprecipitated with cmyc-Ab and western blot was developed through myc antibody. The concentration of CIPK6 expression is shown through protein band intensity below the blot with respect to loading control Rubisco. **(D)** Bacterial count in the col-0, *wrky48, cipk6*, *35S::CIPK6-cMYC*, *35S::CIPK6-cMYC/wrky48* with ± SD of using 8 biological replicate with significance of ***P ≥0.0001, **P≥0.001 and *P ≥ 0.05. **(E)** *Pst* Induced disease phenotype of the indicated genotypes. Experiment with similar results were obtained twice. **(F)** Immunoblot showing the CIPK6 expression in *35S::CIPK6-myc* and *35S::CIPK6/wrky48* plants, where Col-0 and *wrky48* was used as negative control.

## Discussion

Protein phosphorylation is essential mechanism for plant defence, since it controls the signalling pathways that trigger the immunological response against pathogens (Park et al., 2012a). For instance, Arabidopsis CAMTA3, a transcription factor known to be a negative regulator of plant defense, is phosphorylated by MPK3 and MPK6 after flg22 application. The phosphorylation of CAMTA3 results in its destabilisation and promote its nuclear import (Jiang et al., 2020). WRKY family of transcription factors are among the most well-studied proteins, serving as both positive and negative regulators of plant immune responses (Bai et al., 2018; Sheikh et al., 2021). Their ability to form homo- and heterocomplexes, interact with other proteins, and bind to W-box *cis*-elements makes them versatile regulators of plant immunity (Pandey and Somssich, 2009). For instance, AtWRKY33 plays an important role in MPK-mediated camalexin biosynthesis during fungal pathogen attack, while AtWRKY48 negatively impacts basal resistance by suppressing SA mediated signaling pathway (Mao et al., 2011; Xing et al., 2008). These examples highlight the importance of phosphorylation and the dual role of WRKY TFs as either promoter or repressor of immune responses, depending on the type of pathogen.

Our study demonstrates that WRKY48 negatively regulates PTI by repressing the expression of *CIPK6* in response to *Pseudomonas syringae* DC3000 (*Pst*DC3000) infection and flg22 treatment. This repression occurs through its interaction with MPK3, which phosphorylates WRKY48, enhancing its trans-repression activity. Interestingly, WRKY48-mediated downregulation of CIPK6 is associated with decreased ROS production and heightened vulnerability to PstDC3000, which is consistent with its function as a PTI negative regulator.

Our finding also highlights the complexity of signaling networks in plant immunity, where kinases like MPK3 can modulate both positive and negative regulators depending on the context. Furthermore, ability of WRKY48 to suppress *CIPK6* expression adds a novel layer of regulation, linking MAPK signaling to a Calcium-dependent protein kinases (CDPK/CPK)-independent calcium signaling pathways mediated by CIPK6, which is known to regulate ROS and other defense-associated processes(Sardar et al., 2017). The transcriptional regulation of *CIPK6* by WRKY48, as investigated in this study, highlights a specific and preferential interaction between WRKY48 and the *CIPK6* promoter. Our results underline the significance of WRKY48, among other candidate WRKY transcription factors like WRKY28 and WRKY8 of the same class group (IIc), in transcriptional regulation. WRKY48 is previously identified as a negative regulator of plant immunity and its interaction with the *CIPK6* promoter may play a critical role in modulating PTI and ETI responses. Given that CIPK6 is implicated in immune signaling and ROS generation, WRKY48-mediated transcriptional repression of *CIPK6* expression could be a strategy employed by plants to fine-tune defense responses, and a mechanism to revert back to the ground state once the pathogen is contained. The enhanced accumulation of WRKY48 mRNA during the early hours following *PstDC3000* challenge suggests that WRKY48 is rapidly induced as part of the initial immune response of plant. However, its downregulation after 90 minutes may reflect a tightly regulated mechanism where WRKY48 acts to fine-tune the immune response to prevent overactivation, which can be detrimental to the plant. Interestingly, the wrky48 mutants demonstrated increased resistance to *PstDC3000* compared to Col-0 plants, as evidenced by reduced bacterial growth. However, their resistance was not as strong as that of the *cipk6* mutants, indicating that WRKY48-mediated transcriptional repression is not the only mechanism of CIPK6-mediated negative regulation of immunity against *PstDC3000.* The observation that *Pst*-inducible defense marker genes (*PR1*, *PR2*, *ICS1*) are expressed at higher levels in wrky48 mutants compared to Col-0 plants but remain lower than in cipk6 mutants further corroborates the above hypothesis. We also observed the similar expression patterns of PTI marker genes (*ICS1*, *nHL10*, *FRK1*) following flg22 treatment in wrky48 mutantsas compared to Col-0. These findings align with prior studies demonstrating the involvement of WRKY transcription factors in balancing defense activation and minimizing fitness costs.

The increased ROS levels and enhanced expression of ROS-scavenging markers (*CAT1*, *APX1*) in *wrky48* mutants compared to Col-0 confirm the negative regulatory role of WRKY48 in controlling oxidative burst during PTI. However, the lower ROS levels and weaker expression of ROS-related genes in *wrky48* mutants compared to *cipk6* mutants suggest that CIPK6 exerts a more dominant or direct role in negative regulation of ROS. Phosphorylation of WRKY48 by MPK3 enhanced its trans-repression activity, thus serving as a regulatory checkpoint to ensure that immune responses are appropriately scaled. Also, the findings unravel another layer of regulation of ROS generation in maintaining the delicate balance between immune activation and suppression.

The inability of MPK6 to interact with WRKY48 in vitro and in planta suggests distinct roles for MPK3 and MPK6 in PTI signaling, with MPK3 playing a specific role in regulating WRKY48 activity. The enhanced interaction between MPK3 and WRKY48 in response to *PstDC3000* inoculation highlights the pathogen-inducible nature of this association. This finding highlights the necessity of MPK3 for WRKY48 function during PTI. This observation aligns with previous studies showing that phosphorylation by MAP kinases is a common mechanism to regulate WRKY transcription factors during plant immunity (Ishihama and Yoshioka, 2012). These findings are in line with earlier research on MAPK-mediated phosphorylation of transcription factors, including CAMTA3 which controls downstream immunological responses by altering protein activity and localisation (Jiang et al., 2020). The specificity of MPK3-WRKY48 interaction further established the importance of precise MAPK signaling in coordinating plant defense responses.

Phosphorylation by MAPKs plays a crucial role in regulating plant immune responses by modulating the stability, localization, and activity of target proteins (Park et al., 2012b; Thulasi Devendrakumar et al., 2018; Sun and Zhang, 2022a; Sun and Zhang, 2022b; Zhang et al., 2023). In our study, we examined how MPK3-mediated phosphorylation of WRKY48 influences plant immunity by regulating the transcription and translation of CIPK6, a critical component of the immune response. Our findings support the hypothesis that WRKY48 acts as a negative regulator of *CIPK6*, and its activity is modulated by MPK3 through phosphorylation. The data showed that WRKY48 phosphorylation by MPK3 is triggered during a *Pseudomonas syringae* pv. *tomato* DC3000 (*Pst*DC3000) challenge. Plants overexpressing WRKY48 exhibited reduced *CIPK6* transcript and protein levels, resulting in enhanced resistance to bacterial infection. Conversely, in the absence of MPK3, the phosphorylation and stability of WRKY48 were compromised, leading to increased *CIPK6* expression and decreased plant resistance.

WRKY48 phosphorylation was significantly enhanced in response to a *Pst*DC3000 challenge in wild-type plants, aligning with the observed increase in MPK3 expression and activity during early infection. The *in planta* phosphorylation was further confirmed in *35S::WRKY48*/*mpk3* plants, where phosphorylation levels of WRKY48 were significantly lower and unresponsive to pathogen challenge. Phosphorylation of WRKY48 also appeared to stabilize the protein, as evidenced by its increased accumulation/stability up to three hours post-inoculation. Stabilization of phosphorylated WRKY48 ensures its functional role in suppressing *CIPK6* transcription and, thereby, the translation during the critical early phases of pathogen attack. This observation is consistent with the previous studies demonstrating that MAPK-mediated phosphorylation can stabilize proteins to fine-tune immune signaling (Jiang et al., 2020). The suppression of CIPK6 expression by WRKY48 was evident at both transcriptional and translational levels. Following *Pst*DC3000 challenge, *CIPK6* transcript and protein levels were significantly reduced and the suppression was dependent on MPK3, as the absence of MPK3 delayed the reduction in CIPK6 protein levels. These findings substantiate our hypothesis of MPK3-mediated regulation of *CIPK6* expression.

The observed differences in bacterial loads among the stable plant lines (*35S::CIPK6-cmyc*, *35S::WRKY48-GFP/wrky48*, and *35S::WRKY48*/*mpk3*) further illustrate the functional relevance of MPK3-WRKY48-CIPK6 regulation in plant defense. Overexpression of WRKY48 in the *mpk3* mutant background resulted in higher bacterial growth, demonstrating that unphosphorylated WRKY48 cannot effectively repress *CIPK6*. In conclusion, this study highlights the intricate roles of WRKY48 and its phosphorylation by MPK3 in fine-tuning plant immune responses as depicted in the working model (Figure 9). The findings provide new insights into how transcriptional regulators integrate kinase signaling and transcriptional networks to orchestrate defense responses.

**Figure 9.**
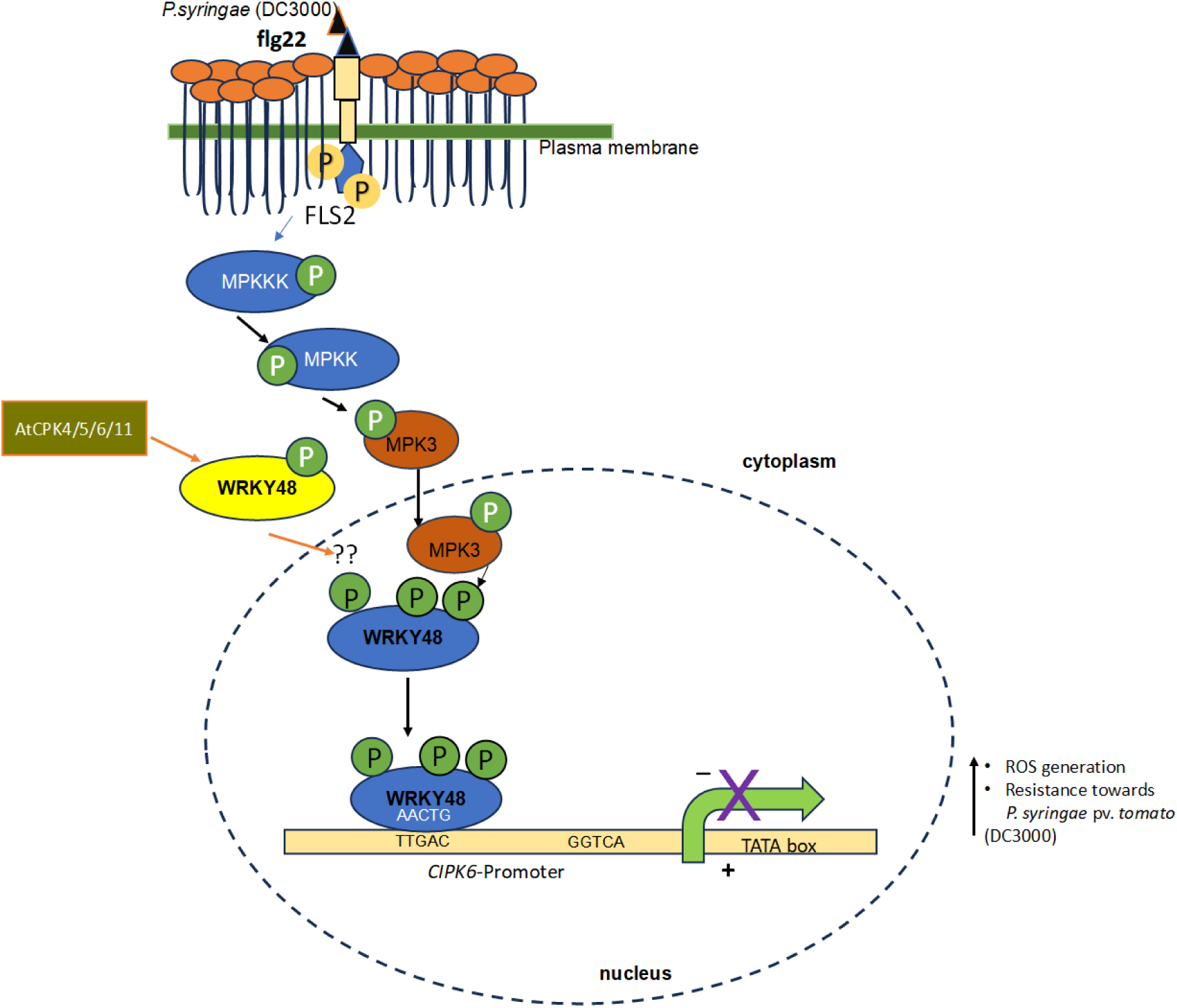
Working model depicting MPK3 mediated phosphorylation of WRKY48 represses CIPK6 Expression during *Pseudomonas syringae* (*Pst*DC3000) infection. The flg22 peptide released by *Pst* DC3000 is recognized by the FLS2 receptor, which initiates a phosphorylation cascade involving MPKKK, MPKK, and MPK3. Once phosphorylated, MPK3 interacts with downstream proteins and transcription factors either in the cytoplasm or within the nucleus. In the nucleus, MPK3 specifically interacts with WRKY48, leading to its phosphorylation. The phosphorylated WRKY48 binds predominantly to the P1 sites of the *CIPK6* promoter, repressing both the *CIPK6* transcript and its protein expression. This repression of *CIPK6* enhances reactive oxygen species (ROS) production, thereby boosting plant resistance.

## Materials and Methods

### Plant materials and growth conditions

All the Arabidopsis genotypes are in Col-0 ecotypes.The T-DNA insertion mutants *wrky48* (salk_0066438), *mpk3* (salk_151594) and *cipk6* (GK-448C12-CS442948) mutants described earlier (Sardar et al., 2017) (Sardar et al, 2017) obtained from ABRC seed stock centre. WRKY48 (*35S::WRKY48-GFP*) overexpression, *mpk3/35S::WRKY48* plants and 35S::CIPK6-cMYC/wrky48 were generated in *wrky48* mutant, and *mpk3/35S::WRKY48* plants were developed in mpk3 background. The pCAMBIA1302 plasmid vector that have hygromycin resistance marker selection for plant selection utilised for *35S::WRKY48-GFP* and *mpk3/35S::WRKY48-GFP* whereas pGWB420 plasmid vector was used for *35S::CIPK6/wrky48* plants, pGWB420 has kanamycin resistance selection marker for plant selection. Arabidopsis plants were grown in controlled growth chamber at 22^0^C, 60-70% relative humidity, and with short day condition (16hrs dark and 8 hours light cycle).

### *Pst*DC3000 inoculation, its count assay and flg22 treatment

Overnight-grown *Pst*DC3000 cultures were suspended in 10 mM MgCl2 and pressure infiltrated in the abaxial side of leaves with a needle-less syringe (Swain et al., 2011). After inoculation plants were covered overnight with plastic dome. The CFU count has been done 3 days post inoculation. Only 10mM MgCl2 was used as mock control wherever required. Leaves tissue for RNA extraction or protein extraction was harvested at the indicated time points. For flg22 treatment, 2uM flg22 prepared in sterile water, was pressure infiltrated in the abaxial side of the leaves. Leaves tissue was harvested at the indicated time point.

### RNA extraction and qPCR analysis

Total RNA from Arabidopsis leaves were isolated by Trizol (Sigma) method. DNase treatment was done to remove DNA before making cDNA. cDNA was made from 1 µg of RNA by Thermo Verso cDNA synthesis kit. The qPCR was done by using 50-100 ng of cDNA and power SYBR green master mix (Applied Biosystem, USA) in ABI-PRISM 7500 FAST sequence detector. For qPCR analysis, typically each sample contained three biological replicates. qPCR was carried out by taking two technical replicates of each biological cDNA. The primers used in the expression analysis are enlisted in supplementary table S1.

### Quantitative measurement of H_2_O_2_ through ROS assay and detection through DAB staining

Hydrogen peroxide accumulation was detected as described earlier (Singh et al., 2018a). In brief, leaves of 4-week-old plants were detached and washed with sterile water. The PstDC3000 inoculation was done through vaccum infiltration after floating the leaves in *Pst*DC3000 solution (0.01 O.D.). The infiltrated leaves were kept in dark at slow shaking (40-50 rpm) for 2-4 h. *Pst*DC3000 solution was taken out and replaced it with DAB staining solution (1 mg/ml DAB in 10 mM Na_2_HPO_4_ and 0.05% Tween 20). Leaves with DAB solution were kept at slow shaking for another 4-6 hrs in dark. After 4 hrs the DAB staining solution was replaced with destaining solution (30: 10: 10 :: Glycerol : Acetic acid : Ethanol) and leaves were incubated overnight at room temperature and photographed.

ROS quantification assay was performed as described earlier (Sardar et al., 2017) , in brief, 4-week-old leaves were detached from Arabidopsis plants and leaf discs of 1.1m^2^ size were sampled through cork-borer and floated on water in 96 well flat bottom griener plates (white color) and kept in dark overnight. After incubating in dark water was removed and washed the leaves once again with sterile water. Then sterile water was replaced with overnight grown culture of *Pst*DC3000 (0.02 O.D.) and vacuum infiltrate the leaf discs for 5 minutes. After vacuum infiltration the elicitation solution 0.2uM luminol (sigma Aldrich) and 20ug/ml of Horseradish peroxidase (Sigma Aldrich). ROS generation was quantified in vivo as luminescence, using a POLARstar Omega (BMG Labtech,UK) 96-well microplate luminometer every 42 s up to 90 minutes. The values are mean of 12 biological replicates.

### Kinase assay

For the MAP Kinase mediated phosphorylation of WRKY48, bacterially expressed recombinant proteins were purified for *in vitro* kinase assay. MPK3, MPK6 and MPK3 kinase dead protein were expressed in BL21DE3-codonPlus, via their cloning in pGEX4T2 and induced with 0.5 mM IPTG. Proteins were purified using Glutathione-Sepharose beads. WRKY48, WRKY28, and WRKY8 were expressed in pET28a^+^ and purified through Ni-NTA beads. Auto-phosphorylation activity of MPK3/6 was checked where MPK3/6 was used as both enzyme and substrate in a kinase buffer (25mM TrisCl pH-7.5, 20mM MgCl_2_, 5mM MnCl_2_ 1mM DTT, 25mM ATP/γ-32P ATP, Bg-phospho 1mM, 1mM Na_3_VO_4_, 2mMEGTA) incubated at 30°C for 30 min, and analyzed by SDS-PAGE. For transphosphorylation activity of MPK3/6, WRKY48 fused with 6X-HIS tag was used as substrate and MPK3 fused with GST was used as kinase for kinase assay.

### Y2H assay

Full length WRKY48 coding sequence was expressed in bait expression plasmid vector pGBKT7 and MPK3 and MPK6 was expressed in pGADT7. Yeast-two-hybrid Gold (Y2H Gold) yeast strain was transformed with both WRKY48-pGBKT7 and MPK3-pGADT7/MPK3-pGADT7. The yeast transformants were selected on synthetic defined medium lacking Leu, Trp. Further to confirm the activation of reporters positive transformants that were already selected on DDO, screened on the synthetic medium lacking leu, Trp, Ade, and histidine (QDO). To confirm the interaction on highly stringent synthetic medium containing 3-AT (Adenine activation) and x-ɑgal (GAL4 activation) in the QDO medium the yeast transformants were serially diluted and allowed to grow by dropping the equal number of cells (10^8^, 10^7^, 10^6^, 10^5^, and 10^4^ cells/ml) for each combination on the above medium. The combination of pGADT7-T7 and pGBKT7-p53 was used as positive control whereas pGADT7-T7 and pGBKT7-lam was used as negative control or pGADT7 only+ pGBKT7-WRKY48 or pKBKT7only + pGADT7-MPK3/6 was used as negative control.

### Bimolecular fluorescence complementation (BiFC)

For performing the BiFC assay, full length coding sequence of MPK3 and MPK6 was amplified and fused with N-terminal YFP fragment of CD3-1648 (pSITE-nEYFP-C1) plasmid vector to generate MPK3-nEYFP or MPK6-nEYFP fusions and WRKY48 CDS was amplified and fused with C-terminal YFP fragment in CD3-1651 (pSITE-cEYFP-N1) plasmid vector to make WRKY48-cEFP fusion construct. The constructed plasmids were transformed in Agrobacterium tumefaciens and transiently expressed in *Nicotiana benthamiana*. The fluorescence was observed after 48h of infiltration under confocal microscope at the same focus and exposure of laser.

### Co-IP and IP

Co-Immunoprecipitation and Immunoprecipitation was done as described by Zhou et al., 2014 and Singh et al., 2018, with some modifications (Zhou et al., 2014). In brief, approximately 250mg of leaves tissue was frozen in liquid N_2_ and total protein was extracted by homogenising the tissue in plant protein extraction buffer (50mM Tris-Cl, pH-8.0, 150mM NaCl, 10% v/v glycerol, 1% v/v Nonidet-40, 0.5% w/v sodium deoxycholate, and plant protease inhibitor (Sigma) 5ul/1ml extraction buffer). 250µg of total protein was incubated with 1µl of required antibody (MPK3 or GFP) for 2 hours at 4^0^C on end-to-end shaker at 13 rpm, then 50µl of protein-A Agarose beads (Invitrogen) was added and kept it on end to end shaking at 4^0^C for 6 hours with same rpm (Singh et al., 2018b). The beads were collected and washed three times with 1X TBST, pH-7.5, beads bound with protein were loaded on 10% SDS-polyacrylamide gels and analysed through immunoblotting through using either GFP (ab290-Abcam), MPK3 (M8318-sigma Aldrich), c-myc (C3956-sigma aldrich), or GST (G7781-sigma aldrich) antibodies.

### EMSA and CHIP

EMSA experiments were performed as described earlier (Giri et al., 2017). In brief, we synthesised 40bps fragment of sense and antisense oligos containing P1 or P2 nucleotides either WT or mutated WRKY binding sites on CIPK6 promoter. Also sense and antisense oligos for 100bps (containing P1 and P2 binding site). The respective sense and antisense oligonucleotides were heated to 100°C for 5 minutes and then gradually cooled to room temperature to obtain an annealed double-stranded 40 bp /100bps DNA fragment . The quantity of protein used for binding with the radiolabelled was 1ug, 2ug, and 4 ug in Figure 1B. in other EMSA experiments we preferred to use 2ug of protein. The samples were resolved on a 6% polyacrylamide gel and shifting was detected through autoradiography (Typhoon-Imager The ChIP experiment was conducted following the protocol described by Saleh et al., 2008 (Saleh et al., 2008). Four-week-old 35S::WRKY48-GFP plants were pressure infiltrated with *Pst* DC3000 suspended in 10 mM MgCl₂ at an optical density (O.D.) of 0.0005 (5x10^5^ CFU/ml), while 10 mM MgCl₂ alone was used for mock infiltration (Singh et al., 2023). Leaf tissue was harvested 24 hours post-infiltration, and chromatin immunoprecipitation was performed using an anti-GFP antibody. The relative abundance of the P1 and P2 elements was quantified via qPCR using primers listed in Table S2.

### Dual-Luciferase reporter assay

The *pCIPK6::LUC* reporter construct was generated by amplifying the 1.2Kb upstream to transcription start site of CIPK6 gene from Arabidopsis gDNA and cloned in pENTR-D-TOPO entry vector followed by mobilisation in p635nfRRF gateway-destination vector. The effector construct was generated in pCAMBIA-1302 by making the fusion of WRKY48-GFP and for MPK3-HA it was generated in pGWB14 gateway-destination vector. The *Agrobacterium* cells were transformed with effector and reporter constructs in combinations and transiently expressed in the same leaf of *Nicotiana benthamiana* at 2 or three different places as shown in the respective figures. Leaf discs from infiltrated *Nicotiana* leaves were harvested at 48 hpi. The discs were homogenized in 1× passive lysis buffer (PLB) provided in the Dual-Luciferase Reporter Assay System (Promega, USA). Firefly luciferase and Renilla luciferase activities were measured using a POLARstar Omega multimode plate reader (BMG Labtech, Germany). Relative luciferase activity (LUC) was determined by normalizing Firefly luciferase activity to Renilla luciferase activity (LUC/REN). The expression of Effector constructs in Nicotiana leaf discs was checked through western blot analysis.

## Acknowledgements

The authors acknowledge the support provided by the central instrumentation facilities and DISC at NIPGR for their assistance in the experiments. They also express their gratitude to the DBT-eLibrary Consortium (DeLCON) for granting access to e-resources. The authors also acknowledge the funding resources provided by the National Institute of Plant Genome Research, Department of Biotechnology (DBT), Ministry of Science and Technology, Government of India. DC acknowledges J.C. Bose Fellowship (JCB/2020/000014) from Science and Engineering Research Board, Department of Science and Technology. NS was supported by CSIR-SRA-pool scientist scheme, Award No. 13(9170-A)/2021-Pool by Council of Scientific and Industrial Research, Govt. of India.

## Authors contributions

NS and DC initiated, conceived, designed, and coordinated the research project. NS wrote the manuscript and DC edited it. NS developed plant lines, reagents, performed experiments. NS and DC analyzed the data.

## Data availability

All data supporting the findings of this study are included in the paper and its supplementary materials available online.

## Supplementary data

**Supplementary Figure 1.**
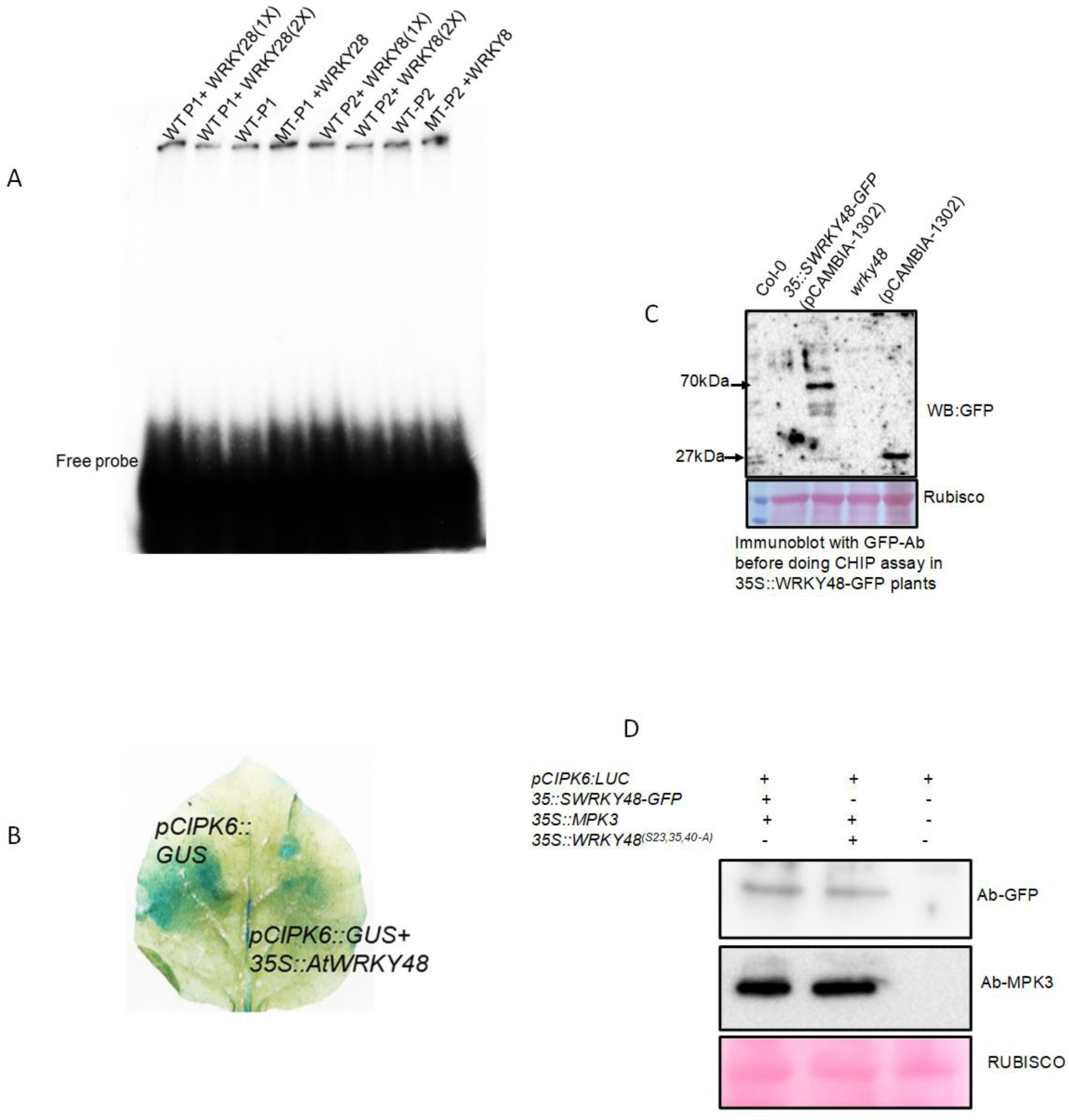
A) EMSA for WRKY28 and WRKY8 with P1 and P2 fragment on *CIPK6* promoter. **B)** Western blot to check the expression of WRKY48 through GFP-antibody in leaves tissue samples collected from pCAMBIA1302-*35S::WRKY48-GFP* and pCAMBIA-*35S-GFP* plants to be used for doing CHIP assay. **C)** Qualitative GUS reporter activity of CIPK6 promoter in the presence of *35S:: WRKY48-GFP*. **D)** Expression of WRKY48, MPK3, and *WRKY48* in the *Nicotiana* leaves tissue used for qualitative and quantitative luciferase-assay, western blot was developed through GFP and MPK3 antibody respectively.

**Supplementary Figure 2.**
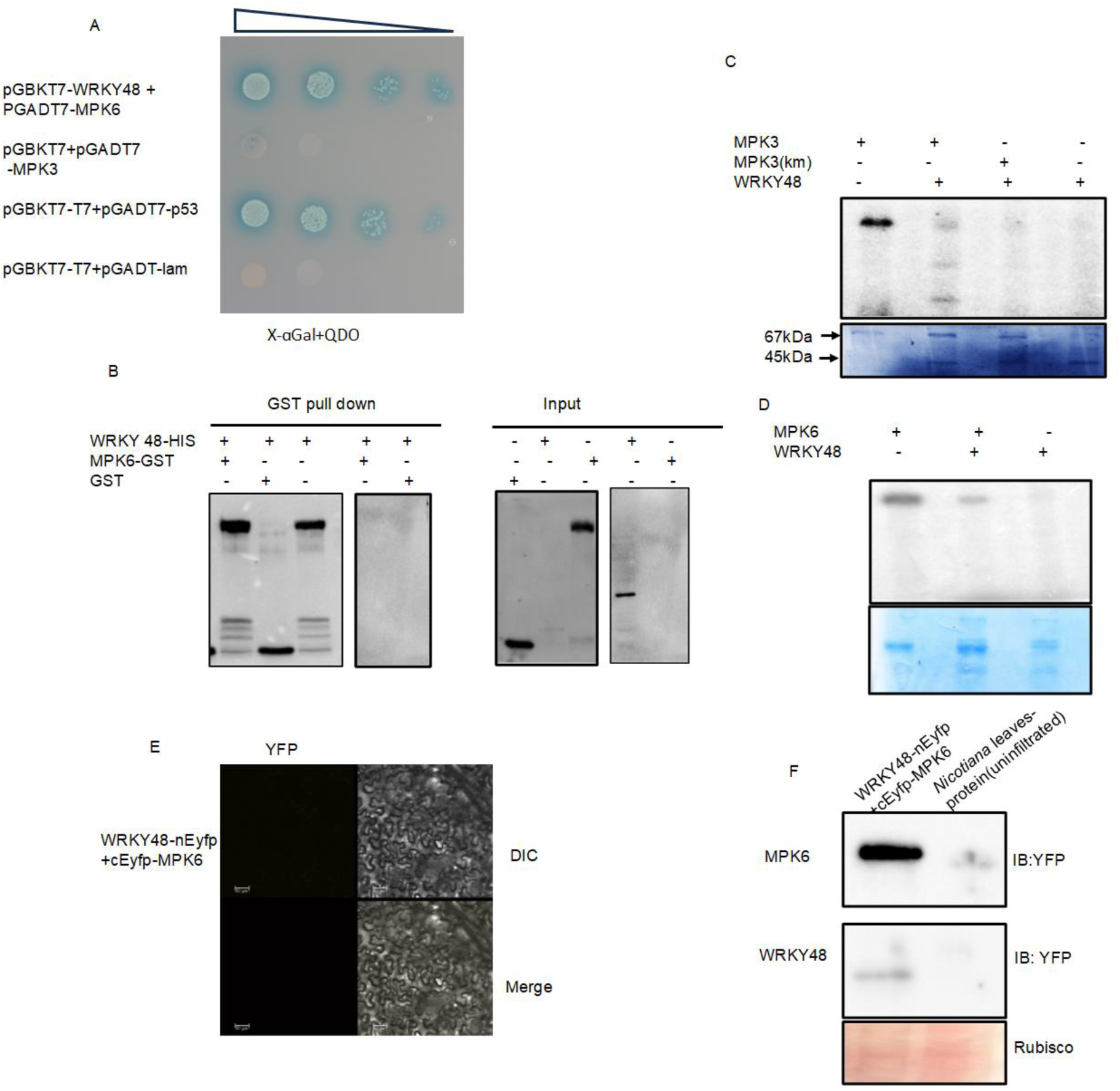
A) Yeast two hybrid assay to check the interaction between MPK6 and WRKY48. **B**) in vitro GST pull down assay to check one to one interaction between MPK6 and WRKY48. **C) and D)** in vitro kinase assay between MPK3-GST and WKY48-HIS and MPK6-GST and WRKY48-HIS. **E)** BIFC assay to check the interaction of WRKY48-nEYFP and MPK6-cEYFP in *Nicotiana benthamiana*. **F)** Immunoblot to check the expression in *Nicotiana* leaves infiltrated with WRKY48-nEYFB+cEYP-MPK6, blots were developed with YFP Ab

**Supplementary Figure 3.**
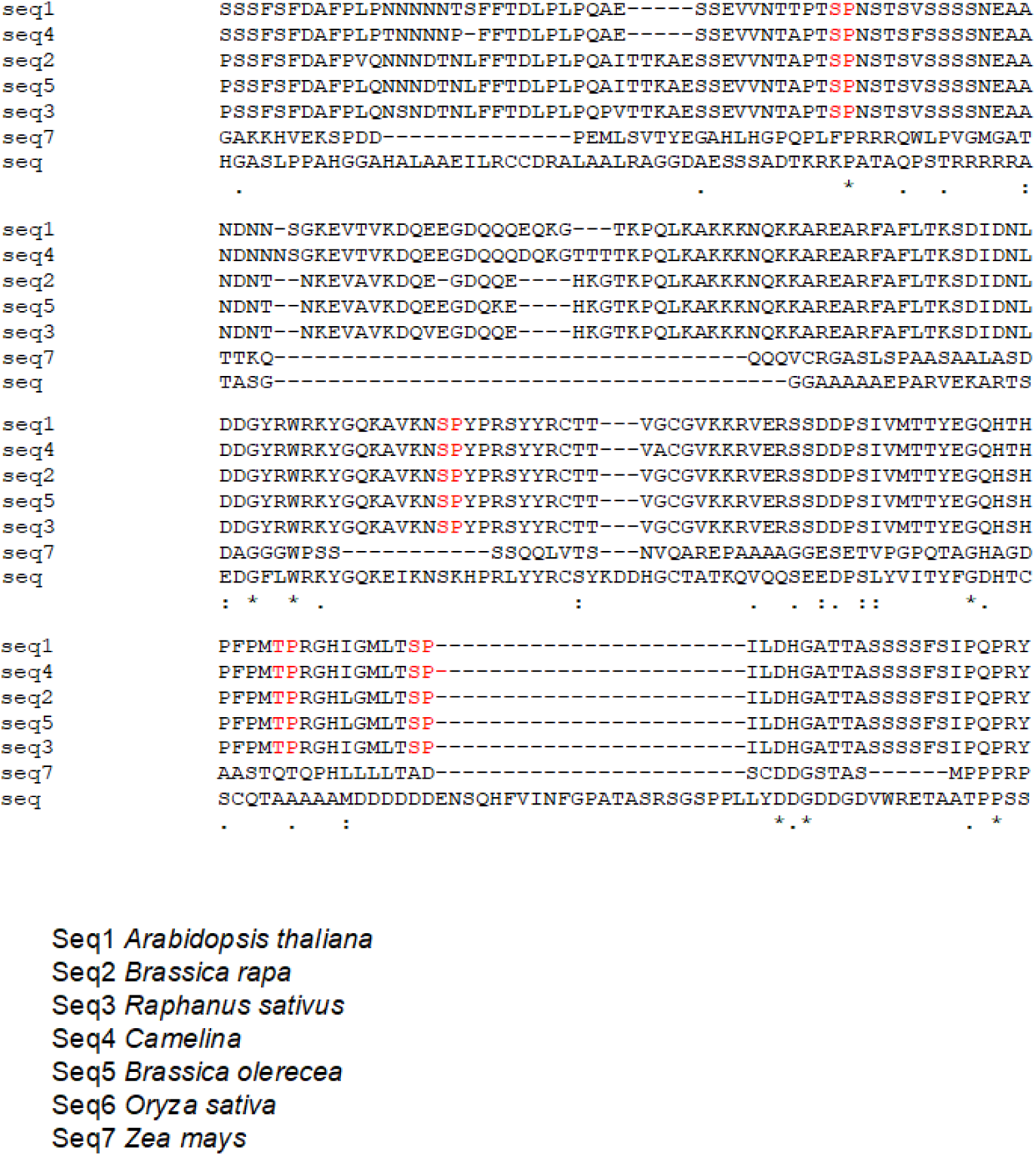
Sequence alignment of WRKY48 from dicot and monocot.

**Supplementary Figure 4.**
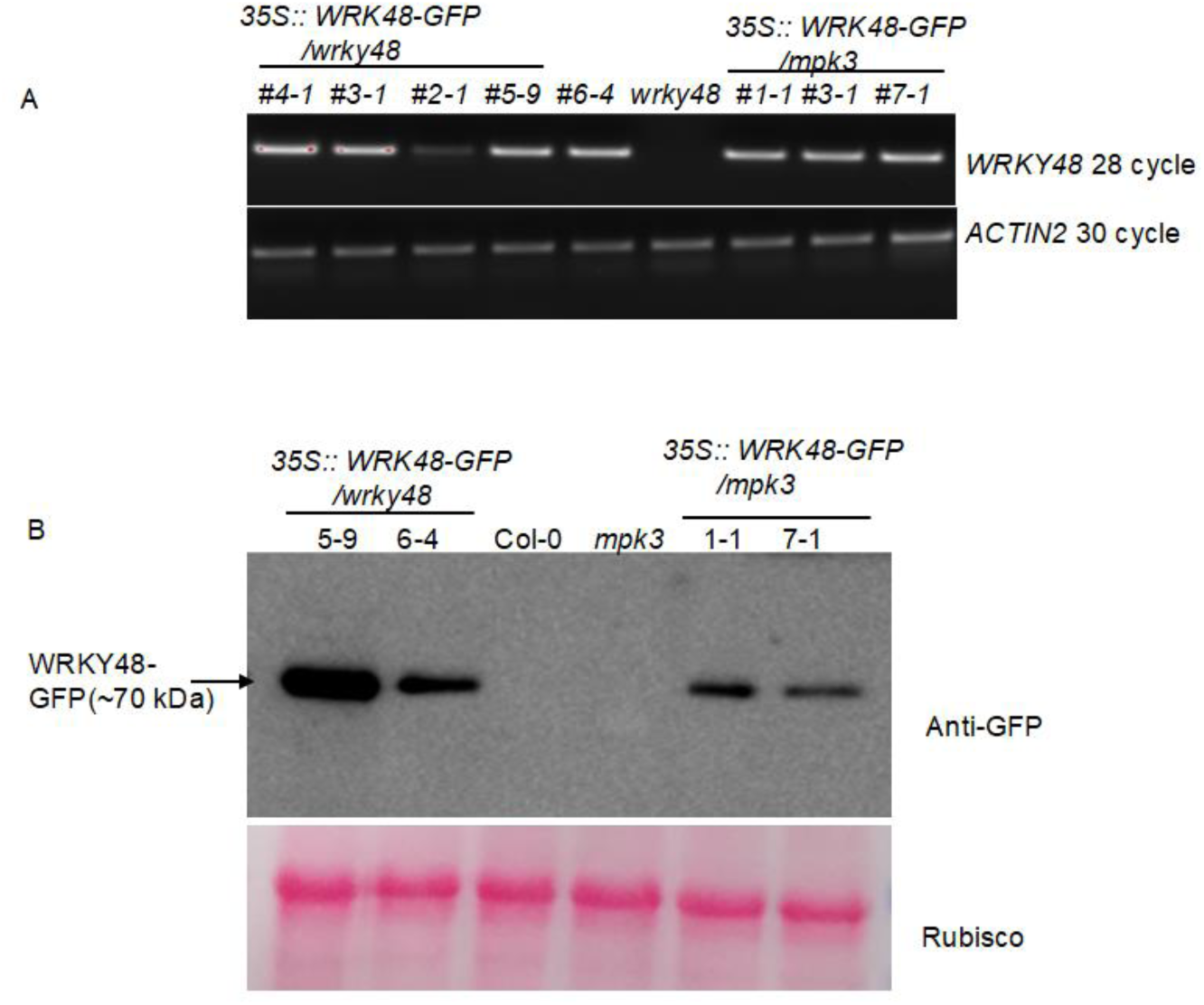
A) Semi-qPCR to confirm the *35S::WRKY48/wrky48* and 35S::WRKY48/*mpk3* homozygous plants in T2 generation B) Western blot to show the WRKY48 expression levels in *35S::WRKY48/wrky48* and 35S::WRKY48/*mpk3* homozygous plants (homozygous line number indicated above the blot).

**Supplementary Figure 5.**
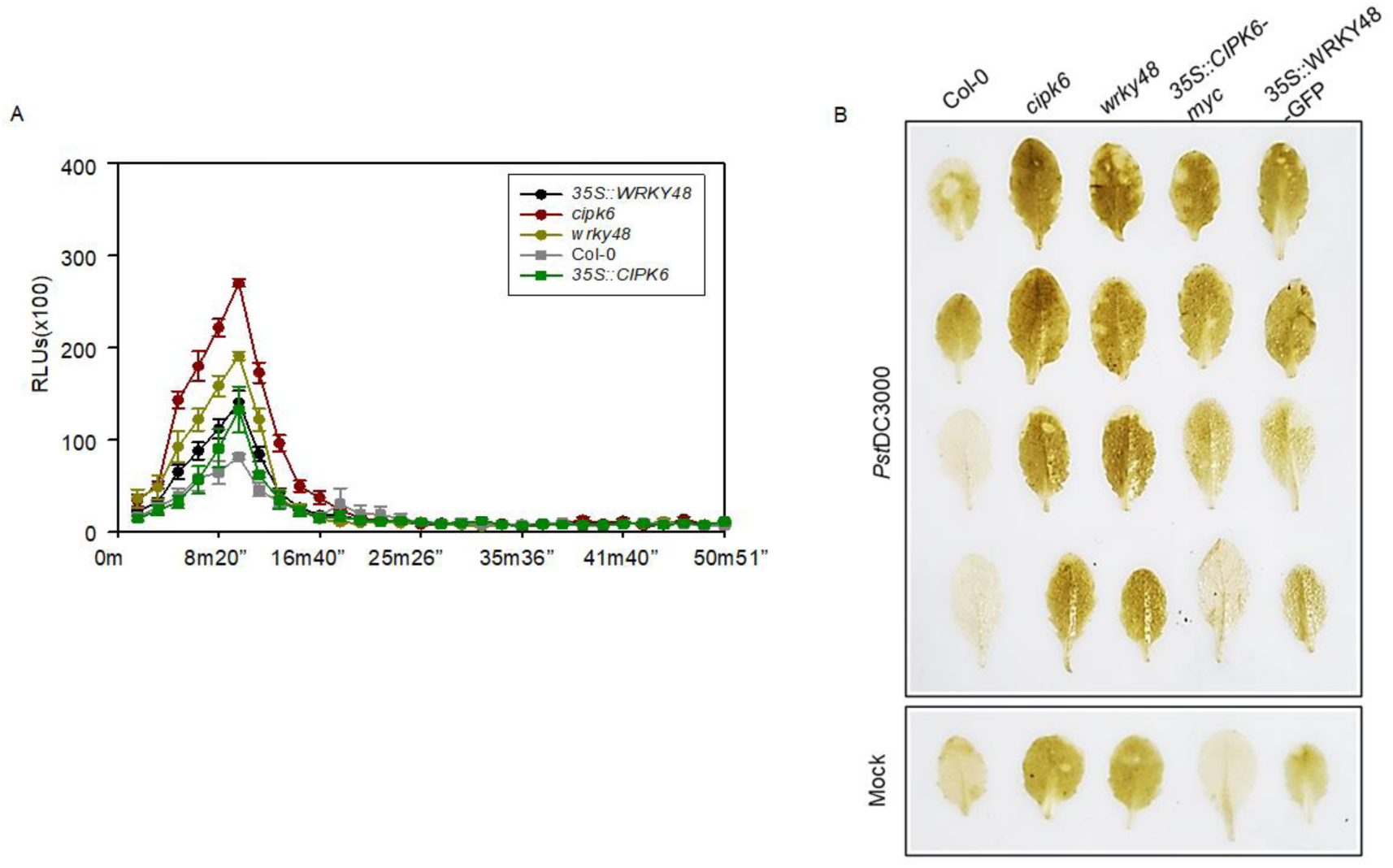
A) ROS kinetics during the indicated time course. Leaf disc of Col-0, *cipk6*, *wrky48*, *35S::CIPK6-myc*, and *35S::WRKY48::GFP* were kept overnight in the water. *Pst*DC3000 (O.D.-0.1) were vacuum infiltrated in the leaf discs and HRP and Luminol at the mentioned concentration in the material methods, were added to run quantify the ROS assay. Data shows ±SE of means where n=12 leaf discs **B)** qualitative DAB staining in Col-0, *cipk6*, *wrky48*, *35S::CIPK6-myc/cipk6*, and *35S::WRKY48-GFP/wrky48*.

**table S1.**
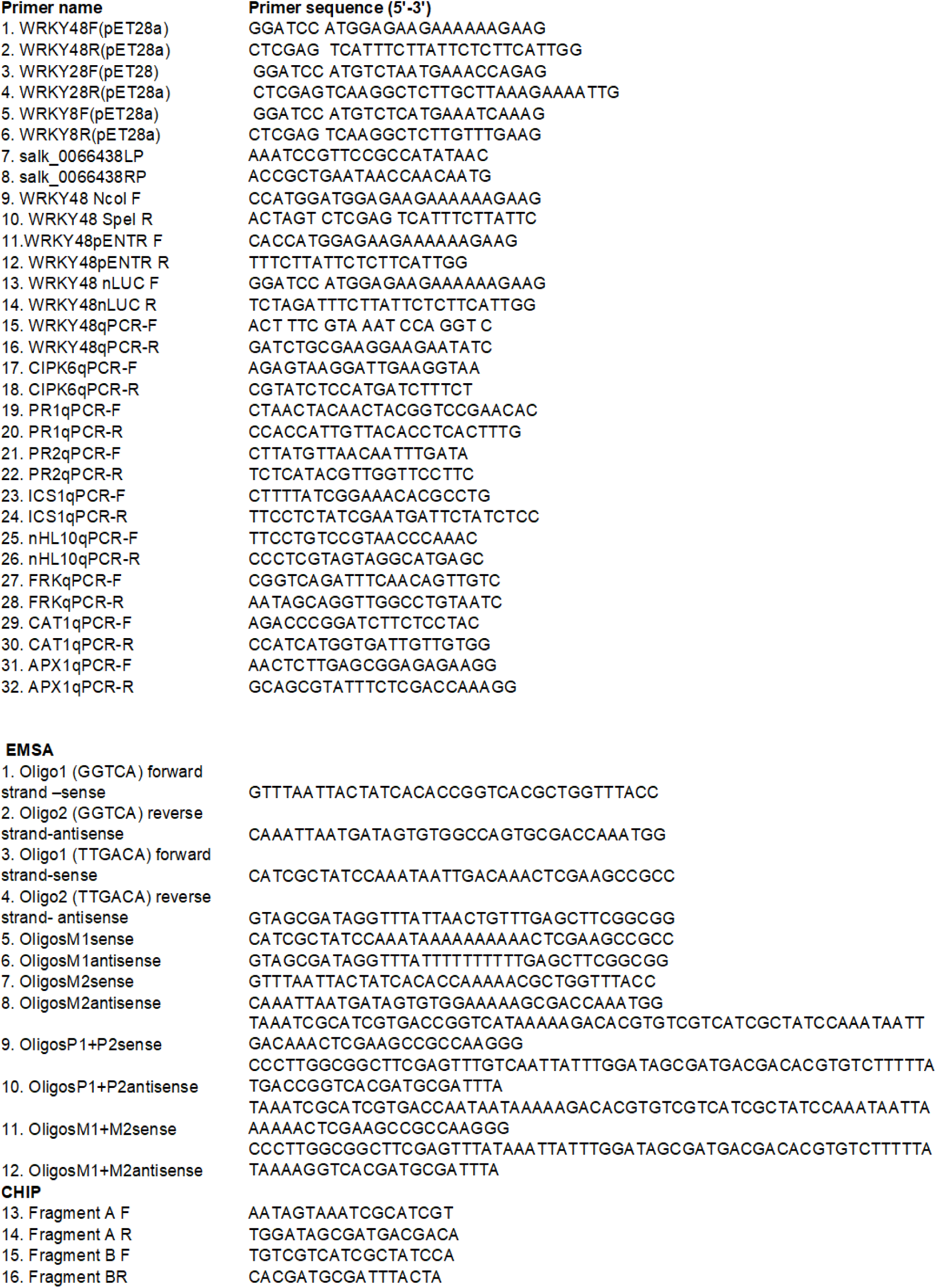
List of Primers used in the study.

## References

Aerts, N., Pereira Mendes, M., and Van Wees, S. C. M. (2021). Multiple levels of crosstalk in hormone networks regulating plant defense. Plant Journal 105:489–504.

Bai, Y., Sunarti, S., Kissoudis, C., Visser, R. G. F., and van der Linden, C. G. (2018). The role of tomato WRKY genes in plant responses to combined abiotic and biotic stresses. Front Plant Sci 9.

Bakshi, M., and Oelmüller, R. (2014). Wrky transcription factors jack of many trades in plants. Plant Signal Behav 9.

Bentham, A. R., de la Concepcion, J. C., Mukhi, N., Zdrzałek, R., Draeger, M., Gorenkin, D., Hughes, R. K., and Banfield, M. J. (2020). A molecular roadmap to the plant immune system. Journal of Biological Chemistry 295:14916–14935.

Bürger, M., and Chory, J. (2019). Stressed Out About Hormones: How Plants Orchestrate Immunity. Cell Host Microbe 26:163–172.

Chen, L., Zhang, L., and Yu, D. (2010). Wounding-Induced WRKY8 Is Involved in Basal Defense in Arabidopsis. Mol Plant Microbe Interact 558:558–565.

Chi, Y., Yang, Y., Zhou, Y., Zhou, J., Fan, B., Yu, J. Q., and Chen, Z. (2013). Protein-protein interactions in the regulation of WRKY transcription factors. Mol Plant 6:287–300.

Deslandes, L., Olivier, J., Dé Ric Theuliè Res†, F., Hirsch, J., Feng, D. X., Bittner-Eddy, P., Beynon, J., and Marco, Y. Resistance to Ralstonia solanacearum in Arabidopsis thaliana is conferred by the recessive RRS1-R gene, a member of a novel family of resistance genes.

Dumanović, J., Nepovimova, E., Natić, M., Kuča, K., and Jaćević, V. (2021). The Significance of Reactive Oxygen Species and Antioxidant Defense System in Plants: A Concise Overview. Front Plant Sci 11.

Eulgem, T., and Somssich, I. E. (2007). Networks of WRKY transcription factors in defense signaling. Curr Opin Plant Biol 10:366–371.

Eulgem, T., Rushton, P. J., Robatzek, S., and Somssich, I. E. (2000). *The WRKY superfamily of plant transcription factors*.

Fernández-Calvo, P., Chini, A., Fernández-Barbero, G., Chico, J. M., Gimenez-Ibanez, S., Geerinck, J., Eeckhout, D., Schweizer, F., Godoy, M., Franco-Zorrilla, J. M., et al. (2011). The Arabidopsis bHLH transcription factors MYC3 and MYC4 are targets of JAZ repressors and act additively with MYC2 in the activation of jasmonate responses. Plant Cell 23:701– 715.

Friso, G., and Van Wijk, K. J. (2015). Posttranslational protein modifications in plant metabolism. Plant Physiol 169:1469–1487.

Gao, Q. M., Venugopal, S., Navarre, D., and Kachroo, A. (2011). Low oleic acid-derived repression of jasmonic acid-inducible defense responses requires the WRKY50 and WRKY51 proteins. Plant Physiol 155:464–476.

Gao, X., Chen, X., Lin, W., Chen, S., Lu, D., Niu, Y., Li, L., Cheng, C., McCormack, M., Sheen, J., et al. (2013). Bifurcation of Arabidopsis NLR Immune Signaling via Ca2+-Dependent Protein Kinases. PLoS Pathog 9.

Giri, M. K., Singh, N., Banday, Z. Z., Singh, V., Ram, H., Singh, D., Chattopadhyay, S., and Nandi, A. K. (2017). GBF1 differentially regulates CAT2 and PAD4 transcription to promote pathogen defense in Arabidopsis thaliana. Plant Journal 91:802–815.

Gorczyca, M., Białas, W., Nicaud, J. M., and Celińska, E. (2024). ‘Mother(Nature) knows best’ – hijacking nature-designed transcriptional programs for enhancing stress resistance and protein production in Yarrowia lipolytica; presentation of YaliFunTome database. Microb Cell Fact 23.

Grunewald, W., Karimi, M., Wieczorek, K., Van De Cappelle, E., Wischnitzki, E., Grundler, F., Inzé, D., Beeckman, T., and Gheysen, G. (2008). A role for AtWRKY23 in feeding site establishment of plant-parasitic nematodes. Plant Physiol 148:358–368.

Higashi, K., Ishiga, Y., Inagaki, Y., Toyoda, K., Shiraishi, T., and Ichinose, Y. (2008). Modulation of defense signal transduction by flagellin-induced WRKY41 transcription factor in Arabidopsis thaliana. Molecular Genetics and Genomics 279:303–312.

Jiang, X., Hoehenwarter, W., Scheel, D., and Lee, J. (2020). Phosphorylation of the CAMTA3 transcription factor triggers its destabilization and nuclear export. Plant Physiol 184:1056– 1071.

Journot-Catalino, H., Somssich, I. E., Roby, D., and Kroj, T. (2006). The transcription factors WRKY11 and WRKY17 act as negative regulators of basal resistance in Arabidopsis thaliana. Plant Cell 18:3289–3302.

Kim, K. C., Lai, Z., Fan, B., and Chen, Z. (2008). Arabidopsis WRKY38 and WRKY62 transcription factors interact with histone deacetylase 19 in basal defense. Plant Cell 20:2357–2371.

Kim, Y., Gilmour, S. J., Chao, L., Park, S., and Thomashow, M. F. (2020). Arabidopsis CAMTA Transcription Factors Regulate Pipecolic Acid Biosynthesis and Priming of Immunity Genes. Mol Plant 13:157–168.

Lai, Z., Vinod, K., Zheng, Z., Fan, B., and Chen, Z. (2008). Roles of Arabidopsis WRKY3 and WRKY4 transcription factors in plant responses to pathogens. BMC Plant Biol 8.

Lee, D. H., Lal, N. K., Lin, Z. J. D., Ma, S., Liu, J., Castro, B., Toruño, T., Dinesh-Kumar, S. P., and Coaker, G. (2020). Regulation of reactive oxygen species during plant immunity through phosphorylation and ubiquitination of RBOHD. Nat Commun 11.

Li, B., Meng, X., Shan, L., and He, P. (2016). Transcriptional Regulation of Pattern-Triggered Immunity in Plants. Cell Host Microbe 19:641–650.

Mao, G., Meng, X., Liu, Y., Zheng, Z., Chen, Z., and Zhang, S. (2011). Phosphorylation of a WRKY transcription factor by two pathogen-responsive MAPKs drives phytoalexin biosynthesis in Arabidopsis. Plant Cell 23:1639–1653.

Meng, X., and Zhang, S. (2013a). MAPK cascades in plant disease resistance signaling. Annu Rev Phytopathol 51:245–266.

Meng, X., and Zhang, S. (2013b). MAPK cascades in plant disease resistance signaling. Annu Rev Phytopathol 51:245–266.

Mithoe, S. C., and Menke, F. L. (2018). Regulation of pattern recognition receptor signalling by phosphorylation and ubiquitination. Curr Opin Plant Biol 45:162–170.

Moore, J. W., Loake, G. J., and Spoel, S. H. (2011). Transcription dynamics in plant immunity. Plant Cell 23:2809–2820.

Mukhtar, M. S., Deslandes, L., Auriac, M. C., Marco, Y., and Somssich, I. E. (2008). The Arabidopsis transcription factor WRKY27 influences wilt disease symptom development caused by Ralstonia solanacearum. Plant Journal 56:935–947.

Murray, S. L., Ingle, R. A., Petersen, L. N., and Denby, K. J. (2007). Basal Resistance Against Pseudomonas syringae in Arabidopsis Involves WRKY53 and a Protein with Homology to a Nematode Resistance Protein. / 1431 MPMI 20:1431–1438.

Narusaka, M., Shirasu, K., Noutoshi, Y., Kubo, Y., Shiraishi, T., Iwabuchi, M., and Narusaka, Y. (2009). RRS1 and RPS4 provide a dual Resistance-gene system against fungal and bacterial pathogens. Plant Journal 60:218–226.

Newton, I. L. G., Woyke, T., Auchtung, T. A., Dilly, G. F., Dutton, R. J., Fisher, M. C., Fontanez, K. M., Lau, E., Stewart, F. J., Richardson, P. M., et al. (2007). The Calyptogena magnifica chemoautotrophic symbiont genome. Science (1979) 315:998–1000.

Ngou, B. P. M., Jones, J. D. G., and Ding, P. (2022). Plant immune networks. Trends Plant Sci 27:255–273.

Pandey, S. P., and Somssich, I. E. (2009). The role of WRKY transcription factors in plant immunity. Plant Physiol 150:1648–1655.

Park, C. J., Caddell, D. F., and Ronald, P. C. (2012a). Protein phosphorylation in plant immunity: Insights into the regulation of pattern recognition receptor-mediated signaling. Front Plant Sci 3.

Park, C. J., Caddell, D. F., and Ronald, P. C. (2012b). Protein phosphorylation in plant immunity: Insights into the regulation of pattern recognition receptor-mediated signaling. Front Plant Sci 3.

Pearson, G., Robinson, F., Gibson, T. B., Xu, B.-E., Karandikar, M., Berman, K., and Cobb, M. H. (2001). Mitogen-Activated Protein (MAP) Kinase Pathways: Regulation and Physiological Functions*.

Phukan, U. J., Jeena, G. S., and Shukla, R. K. (2016). WRKY transcription factors: Molecular regulation and stress responses in plants. Front Plant Sci 7.

Plotnikov, A., Zehorai, E., Procaccia, S., and Seger, R. (2011). The MAPK cascades: Signaling components, nuclear roles and mechanisms of nuclear translocation. Biochim Biophys Acta Mol Cell Res 1813:1619–1633.

Qia, Z., Verma, R., Gehring, C., Yamaguchi, Y., Zhao, Y., Ryan, C. A., and Berkowitz, G. A. (2010). Ca 2+ signaling by plant Arabidopsis thaliana Pep peptides depends on AtPepR1, a receptor with guanylyl cyclase activity, and cGMP-activated Ca 2+ channels. Proc Natl Acad Sci U S A 107:21193–21198.

Ren, C.-X., Chen, S.-Y., He, Y.-H., Xu, Y.-P., Yang, J., and Cai, X.-Z. (2024). Fine-tuning of the dual-role transcription factor WRKY8 via differential phosphorylation for robust broad-spectrum plant immunity. Plant Commun Advance Access published December 2024, doi:10.1016/j.xplc.2024.101072.

Ribet, D., and Cossart, P. (2010). Pathogen-mediated posttranslational modifications: A re-emerging field. Cell 143:694–702.

Saleh, A., Alvarez-Venegas, R., and Avramova, Z. (2008). An efficient chromatin immunoprecipitation (ChIP) protocol for studying histone modifications in Arabidopsis plants. Nat Protoc 3:1018–1025.

Sardar, A., Nandi, A. K., and Chattopadhyay, D. (2017). CBL-interacting protein kinase 6 negatively regulates immune response to Pseudomonas syringae in Arabidopsis. J Exp Bot 68:3573–3584.

Sheikh, A. H., Fraz Hussain, R. M., Tabassum, N., Badmi, R., Marillonnet, S., Scheel, D., Lee, J., and Sinha, A. (2021). Possible role of WRKY transcription factors in regulating immunity in Oryza sativa ssp. indica. Physiol Mol Plant Pathol 114.

Singh, A., Sagar, S., and Biswas, D. K. (2017). Calcium Dependent Protein Kinase, a Versatile Player in Plant Stress Management and Development. CRC Crit Rev Plant Sci 36:336–352.

Singh, N., Swain, S., Singh, A., and Nandi, A. K. (2018a). AtOZF1 positively regulates defense against bacterial pathogens and NPR1-independent salicylic acid signaling. Molecular Plant-Microbe Interactions 31:323–333.

Singh, N., Swain, S., Singh, A., and Nandi, A. K. (2018b). AtOZF1 positively regulates defense against bacterial pathogens and NPR1-independent salicylic acid signaling. Molecular Plant-Microbe Interactions 31:323–333.

Singh, D., Patil, V., Kumar, R., Gautam, J. K., Singh, V., and Nandi, A. K. (2023). RSI1/FLD and its epigenetic target RRTF1 are essential for the retention of infection memory in Arabidopsis thaliana. Plant Journal 115:662–677.

Sun, T., and Zhang, Y. (2022a). MAP kinase cascades in plant development and immune signaling. EMBO Rep 23.

Sun, T., and Zhang, Y. (2022b). MAP kinase cascades in plant development and immune signaling. EMBO Rep 23.

Swain, S., Roy, S., Shah, J., van Wees, S., Pieterse, C. M., and Nandi, A. K. (2011). Arabidopsis thaliana cdd1 mutant uncouples the constitutive activation of salicylic acid signalling from growth defects. Mol Plant Pathol 12:855–865.

Thordal-Christensen, H. (2020). A holistic view on plant effector-triggered immunity presented as an iceberg model. Cellular and Molecular Life Sciences 77:3963–3976.

Thulasi Devendrakumar, K., Li, X., and Zhang, Y. (2018). MAP kinase signalling: interplays between plant PAMP- and effector-triggered immunity. Cellular and Molecular Life Sciences 75:2981–2989.

Transcription Factors Involved in Plant Resistance to Pathogens.

Tsuda, K., and Somssich, I. E. (2015). Transcriptional networks in plant immunity. New Phytologist 206:932–947.

Waheed, A., Haxim, Y., Islam, W., Kahar, G., Liu, X., and Zhang, D. (2022). Role of pathogen’s effectors in understanding host-pathogen interaction. Biochim Biophys Acta Mol Cell Res 1869.

Wang, D., Amornsiripanitch, N., and Dong, X. (2006). A genomic approach to identify regulatory nodes in the transcriptional network of systemic acquired resistance in plants. PLoS Pathog 2:1042–1050.

Wang, H., Cheng, X., Yin, D., Chen, D., Luo, C., Liu, H., and Huang, C. (2023). Advances in the Research on Plant WRKY Transcription Factors Responsive to External Stresses. Curr Issues Mol Biol 45:2861–2880.

Xing, D. H., Lai, Z. B., Zheng, Z. Y., Vinod, K. M., Fan, B. F., and Chen, Z. X. (2008). Stress- and pathogen-induced Arabidopsis WRKY48 is a transcriptional activator that represses plant basal defense. Mol Plant 1:459–470.

Xu, X., Chen, C., Fan, B., and Chen, Z. (2006). Physical and functional interactions between pathogen-induced Arabidopsis WRKY18, WRKY40, and WRKY60 transcription factors. Plant Cell 18:1310–1326.

Yuan, M., Jiang, Z., Bi, G., Nomura, K., Liu, M., Wang, Y., Cai, B., Zhou, J. M., He, S. Y., and Xin, X. F. (2021). Pattern-recognition receptors are required for NLR-mediated plant immunity. Nature 592:105–109.

Zhang, M., and Zhang, S. (2022). Mitogen-activated protein kinase cascades in plant signaling. J Integr Plant Biol 64:301–341.

Zhang, W. J., Zhou, Y., Zhang, Y., Su, Y. H., and Xu, T. (2023). Protein phosphorylation: A molecular switch in plant signaling. Cell Rep 42.

Zheng, Z., Qamar, S. A., Chen, Z., and Mengiste, T. (2006). Arabidopsis WRKY33 transcription factor is required for resistance to necrotrophic fungal pathogens. Plant Journal 48:592– 605.

Zhou, J., Wu, S., Chen, X., Liu, C., Sheen, J., Shan, L., and He, P. (2014). The Pseudomonas syringae effector HopF2 suppresses Arabidopsis immunity by targeting BAK1. Plant Journal 77:235–245.

