## Supplementary file for "MPK3 mediated phosphorylation of WRKY48 down regulates CIPK6 expression during *Pst*DC3000 challenge in Arabidopsis"

Supplementary Figure 1

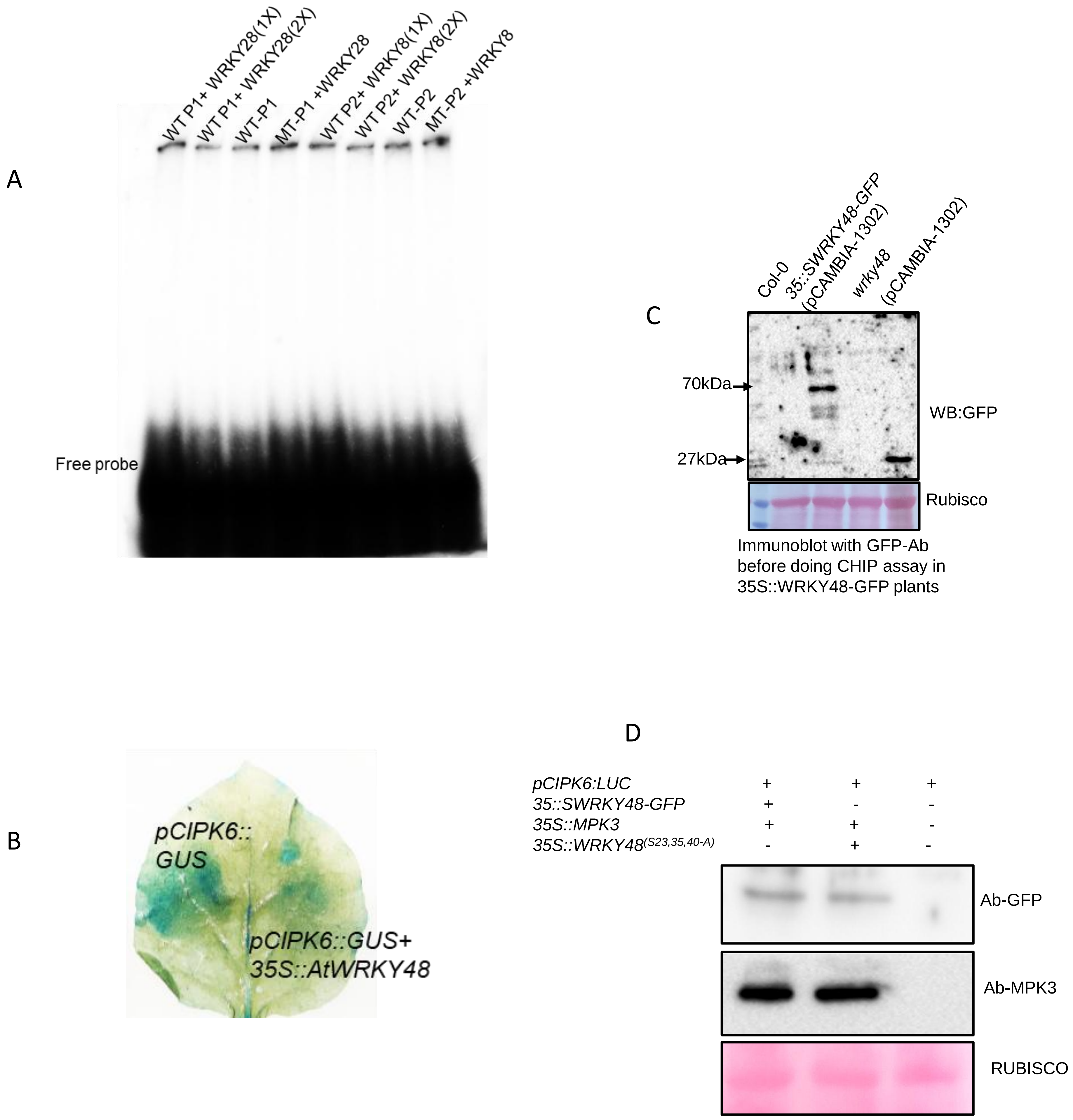


**Supplementary Figure 1. A)** EMSA for WRKY28 and WRKY8 with P1 and P2 fragment on *CIPK6* promoter. **B)** Western blot to check the expression of WRKY48 through GFP-antibody in leaves tissue samples collected from pCAMBIA1302-*35S::WRKY48-GFP* and pCAMBIA-*35S-GFP* plants to be used for doing CHIP assay. **C)** Qualitative GUS reporter activity of *CIPK6* promoter in the presence of *35S:: WRKY48-GFP*. **D)** Expression of WRKY48, MPK3, and *WRKY48* in the *Nicotiana* leaves tissue used for qualitative and quantitative luciferase-assay, western blot was developed through GFP and MPK3 antibody respectively.

Y2H assay WRKY48+MPK6

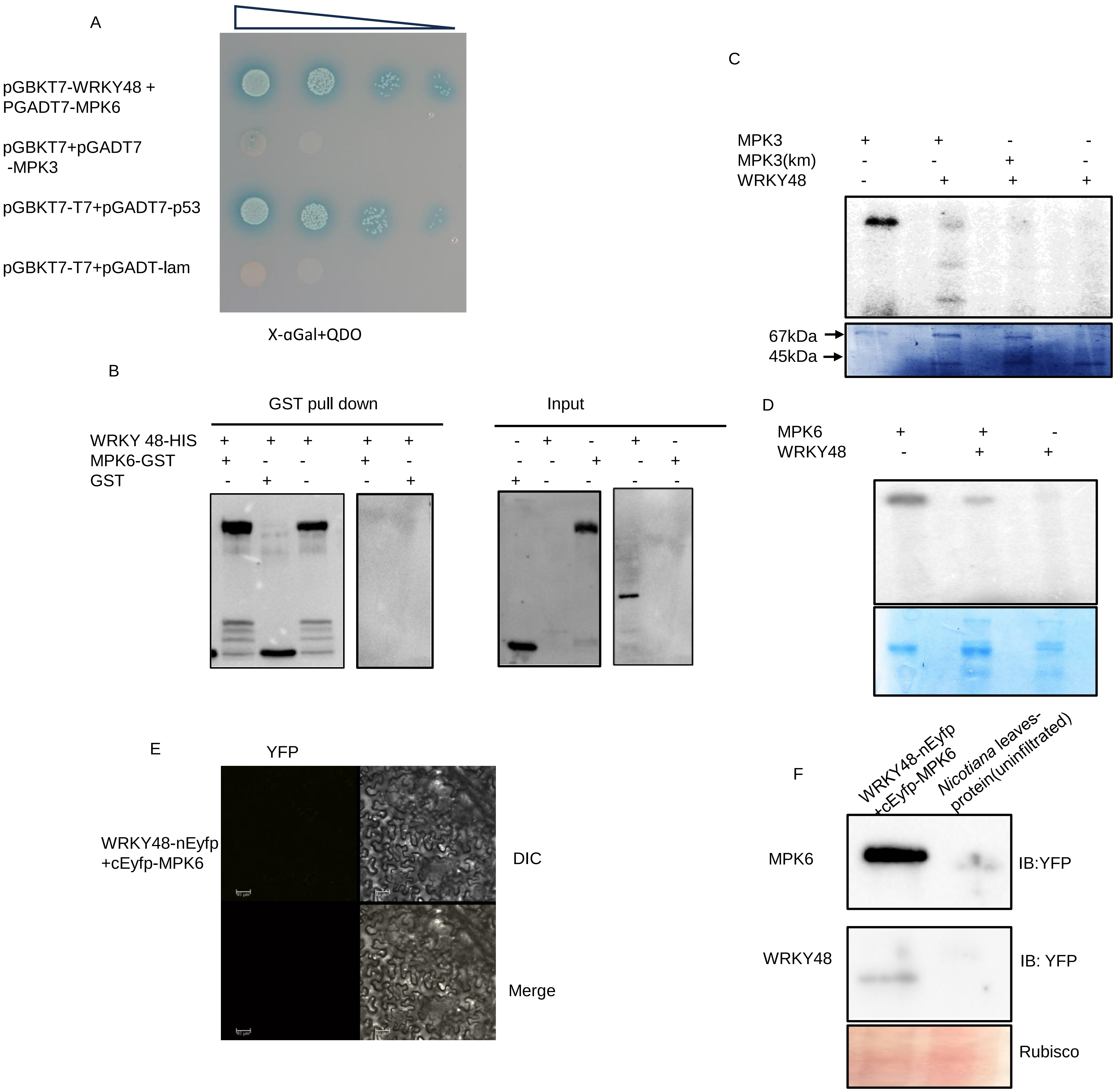


**Supplementary Figure 2. A)** Yeast two hybrid assay to check the interaction between MPK6 and WRKY48. **B**) in vitro GST pull down assay to check one to one interaction between MPK6 and WRKY48. **C) and D)** in vitro kinase assay between MPK3-GST and WKY48-HIS and MPK6-GST and WRKY48-HIS. **E)** BIFC assay to check the interaction of WRKY48-nEYFP and MPK6-cEYFP in *Nicotiana benthamiana*. **F)** Immunoblot to check the expression in *Nicotiana* leaves infiltrated with WRKY48-nEYFB+cEYP-MPK6, blots were developed with YFP Ab.

```
seq1            SSSFSFDAFPLPNNNNNTSFFTDLPLPQAE-----SSEVVNTTPTSPNSTSVSSSSNEAA
seq4            SSSFSFDAFPLPTNNNNP-FFTDLPLPQAE-----SSEVVNTAPTSPNSTSFSSSSNEAA
seq2            PSSFSFDAFPVQNNNDTNLFFTDLPLPQAITTKAESSEVVNTAPTSPNSTSVSSSSNEAA
seq5            PSSFSFDAFPLQNNNDTNLFFTDLPLPQAITTKAESSEVVNTAPTSPNSTSVSSSSNEAA
seq3            PSSFSFDAFPLQNSNDTNLFFTDLPLPQPVTTKAESSEVVNTAPTSPNSTSVSSSSNEAA
seq7            GAKKHVEKSPDD--------------PEMLSVTYEGAHLHGPQPLFPRRRQWLPVGMGAT
seq             HGASLPPAHGGAHALAAEILRCCDRALAALRAGGDAESSSADTKRKPATAQPSTRRRRRA
                 .                                 .          *   .  .     :

seq1            NDNN-SGKEVTVKDQEEGDQQQEQKG---TKPQLKAKKKNQKKAREARFAFLTKSDIDNL
seq4            NDNNNSGKEVTVKDQEEGDQQQDQKGTTTTKPQLKAKKKNQKKAREARFAFLTKSDIDNL
seq2            NDNT--NKEVAVKDQE-GDQQE----HKGTKPQLKAKKKNQKKAREARFAFLTKSDIDNL
seq5            NDNT--NKEVAVKDQEEGDQKE----HKGTKPQLKAKKKNQKKAREARFAFLTKSDIDNL
seq3            NDNT--NKEVAVKDQVEGDQQE----HKGTKPQLKAKKKNQKKAREARFAFLTKSDIDNL
seq7            TTKQ-----------------------------------QQQVCRGASLSPAASAALASD
seq             TASG--------------------------------------GGAAAAAEPARVEKARTS

seq1            DDGYRWRKYGQKAVKNSPYPRSYYRCTT---VGCGVKKRVERSSDDPSIVMTTYEGQHTH
seq4            DDGYRWRKYGQKAVKNSPYPRSYYRCTT---VACGVKKRVERSSDDPSIVMTTYEGQHTH
seq2            DDGYRWRKYGQKAVKNSPYPRSYYRCTT---VGCGVKKRVERSSDDPSIVMTTYEGQHSH
seq5            DDGYRWRKYGQKAVKNSPYPRSYYRCTT---VGCGVKKRVERSSDDPSIVMTTYEGQHSH
seq3            DDGYRWRKYGQKAVKNSPYPRSYYRCTT---VGCGVKKRVERSSDDPSIVMTTYEGQHSH
seq7            DAGGGWPSS-----------SSQQLVTS---NVQAREPAAAAGGESETVPGPQTAGHAGD
seq             EDGFLWRKYGQKEIKNSKHPRLYYRCSYKDDHGCTATKQVQQSEEDPSLYVITYFGDHTC
                : *  * .                  :            .  . :. ::      *.

seq1            PFPMTPRGHIGMLTSP-----------------------ILDHGATTASSSSFSIPQPRY
seq4            PFPMTPRGHIGMLTSP-----------------------ILDHGATTASSSSFSIPQPRY
seq2            PFPMTPRGHLGMLTSP-----------------------ILDHGATTASSSSFSIPQPRY
seq5            PFPMTPRGHLGMLTSP-----------------------ILDHGATTASSSSFSIPQPRY
seq3            PFPMTPRGHIGMLTSP-----------------------ILDHGATTASSSSFSIPQPRY
seq7            AASTQTQPHLLLLTAD-----------------------SCDDGSTAS------MPPPRP
seq             SCQTAAAAAMDDDDDDENSQHFVINFGPATASRSGSPPLLYDDGDDGDVWRETAATPPSS
                .    .   :                               *.*           . *
```

Seq1 *Arabidopsis thaliana*
Seq2 *Brassica rapa*
Seq3 *Raphanus sativus*
Seq4 *Camelina*
Seq5 *Brassica olerecea*
Seq6 *Oryza sativa*
Seq7 *Zea mays*

**Supplementary Figure 3** Sequence alignment of WRKY48 from dicot and monocot

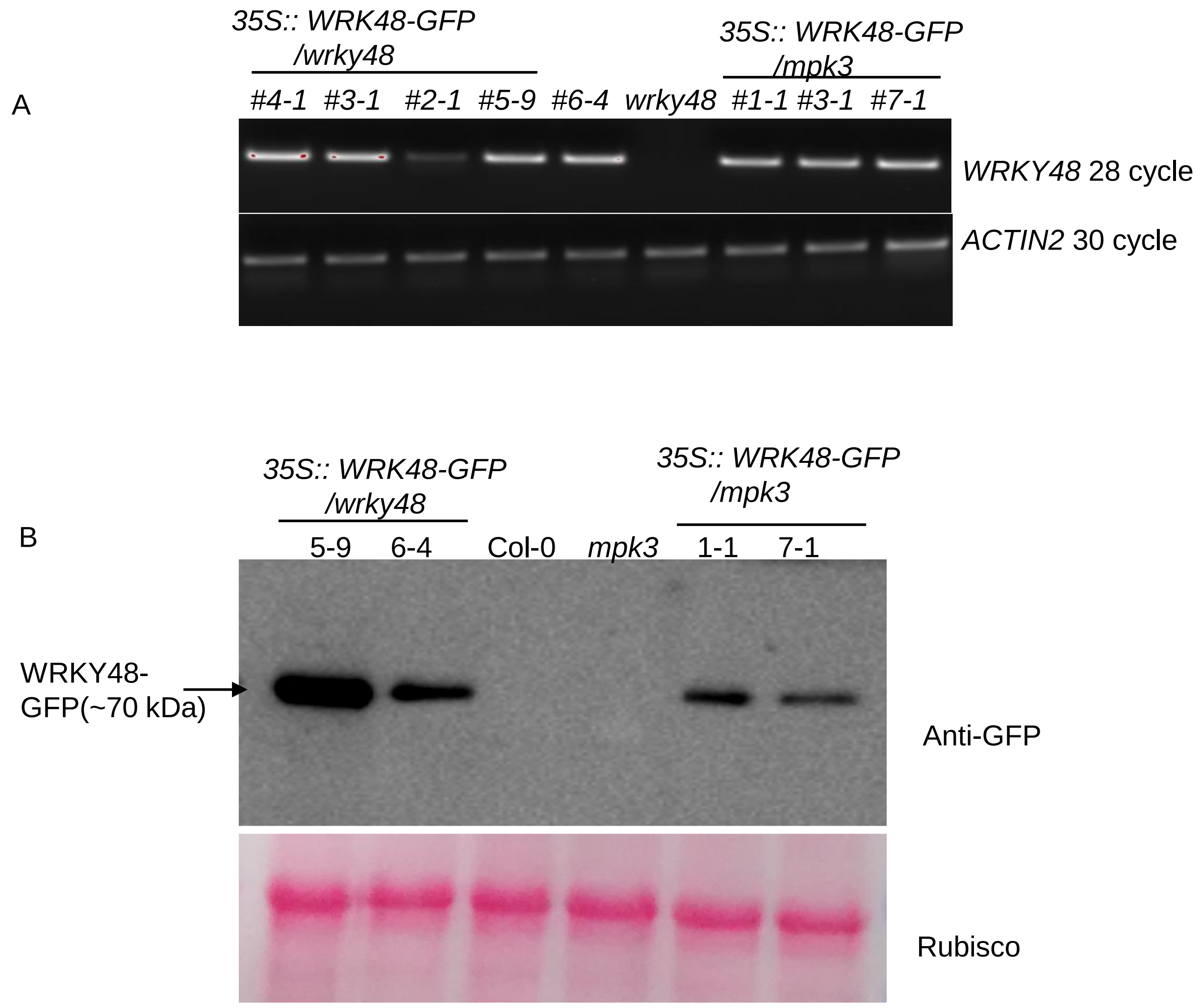


**Supplementary Figure 3. A)** Semi-qPCR to confirm the *35S::WRKY48/wrky48* and 35S::WRKY48/*mpk3* homozygous plants in T2 generation **B)** Western blot to show the WRKY48 expression levels in *35S::WRKY48/wrky48* and 35S::WRKY48/*mpk3* homozygous plants (homozygous line number indicated above the blot).

A

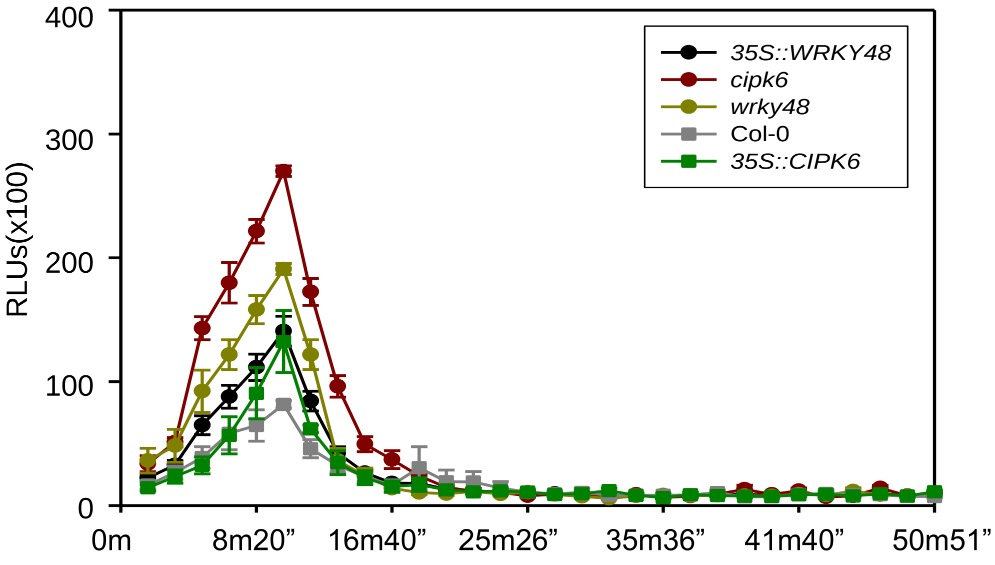


B

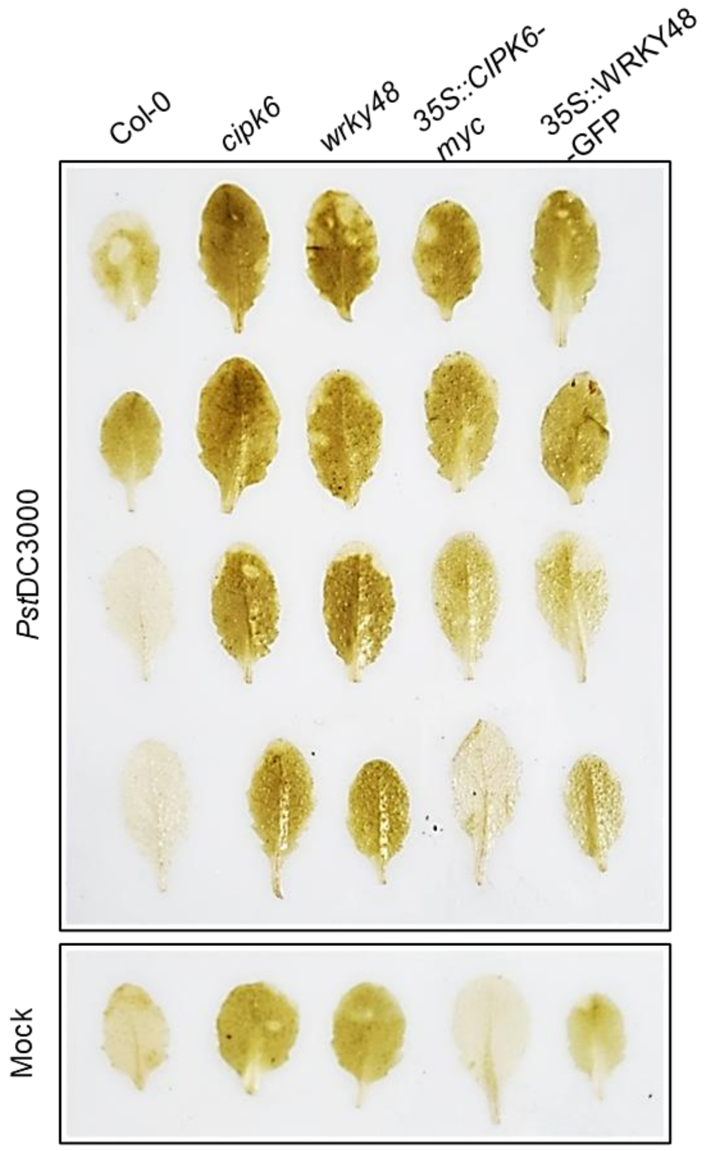


**Supplementary Figure 4. A)** ROS kinetics during the indicated time course. Leaf disc of Col-0, cipk6, wrky48, *35S::CIPK6-myc*, and *35S::WRKY48::GFP* were kept overnight in the water. *Pst*DC3000 (O.D.- 0.1) were vacuum infiltrated in the leaf discs and HRP and Luminol at the mentioned concentration in the material methods, were added to run quantify the ROS assay. Data shows ±SE of means where n=12 leaf discs **B)** qualitative DAB staining in Col-0, *cipk6*, *wrky48*, *35S::CIPK6-myc*, and *35S::WRKY48:-GFP.*

Table S1: Primers list

| **Primer name** | **Primer sequence (5'-3')** |
|---|---|
| 1. WRKY48F(pET28a) | GGATCC ATGGAGAAGAAAAAAGAAG |
| 2. WRKY48R(pET28a) | CTCGAG TCATTTCTTATTCTCTTCATTGG |
| 3. WRKY28F(pET28) | GGATCC ATGTCTAATGAAACCAGAG |
| 4. WRKY28R(pET28a) | CTCGAGTCAAGGCTCTTGCTTAAAGAAAATTG |
| 5. WRKY8F(pET28a) | GGATCC ATGTCTCATGAAATCAAAG |
| 6. WRKY8R(pET28a) | CTCGAG TCAAGGCTCTTGTTTGAAG |
| 7. salk_0066438LP | AAATCCGTTCCGCCATATAAC |
| 8. salk_0066438RP | ACCGCTGAATAACCAACAATG |
| 9. WRKY48 NcoI F | CCATGGATGGAGAAGAAAAAAGAAG |
| 10. WRKY48 SpeI R | ACTAGT CTCGAG TCATTTCTTATTC |
| 11.WRKY48pENTR F | CACCATGGAGAAGAAAAAAGAAG |
| 12. WRKY48pENTR R | TTTCTTATTCTCTTCATTGG |
| 13. WRKY48 nLUC F | GGATCC ATGGAGAAGAAAAAAGAAG |
| 14. WRKY48nLUC R | TCTAGATTTCTTATTCTCTTCATTGG |
| 15. WRKY48qPCR-F | ACT TTC GTA AAT CCA GGT C |
| 16. WRKY48qPCR-R | GATCTGCGAAGGAAGAATATC |
| 17. CIPK6qPCR-F | AGAGTAAGGATTGAAGGTAA |
| 18. CIPK6qPCR-R | CGTATCTCCATGATCTTTCT |
| 19. PR1qPCR-F | CTAACTACAACTACGGTCCGAACAC |
| 20. PR1qPCR-R | CCACCATTGTTACACCTCACTTTG |
| 21. PR2qPCR-F | CTTATGTTAACAATTTGATA |
| 22. PR2qPCR-R | TCTCATACGTTGGTTCCTTC |
| 23. ICS1qPCR-F | CTTTTATCGGAAACACGCCTG |
| 24. ICS1qPCR-R | TTCCTCTATCGAATGATTCTATCTCC |
| 25. nHL10qPCR-F | TTCCTGTCCGTAACCCAAAC |
| 26. nHL10qPCR-R | CCCTCGTAGTAGGCATGAGC |
| 27. FRKqPCR-F | CGGTCAGATTTCAACAGTTGTC |
| 28. FRKqPCR-R | AATAGCAGGTTGGCCTGTAATC |
| 29. CAT1qPCR-F | AGACCCGGATCTTCTCCTAC |
| 30. CAT1qPCR-R | CCATCATGGTGATTGTTGTGG |
| 31. APX1qPCR-F | AACTCTTGAGCGGAGAGAAGG |
| 32. APX1qPCR-R | GCAGCGTATTTCTCGACCAAAGG |
| **EMSA** | |
| 1. Oligo1 (GGTCA) forward strand –sense | GTTTAATTACTATCACACCGGTCACGCTGGTTTACC |
| 2. Oligo2 (GGTCA) reverse strand-antisense | CAAATTAATGATAGTGTGGCCAGTGCGACCAAATGG |
| 3. Oligo1 (TTGACA) forward strand-sense | CATCGCTATCCAAATAATTGACAAACTCGAAGCCGCC |
| 4. Oligo2 (TTGACA) reverse strand- antisense | GTAGCGATAGGTTTATTAACTGTTTGAGCTTCGGCGG |
| 5. OligosM1sense | CATCGCTATCCAAATAAAAAAAAAACTCGAAGCCGCC |
| 6. OligosM1antisense | GTAGCGATAGGTTTATTTTTTTTTTGAGCTTCGGCGG |
| 7. OligosM2sense | GTTTAATTACTATCACACCAAAAACGCTGGTTTACC |
| 8. OligosM2antisense | CAAATTAATGATAGTGTGGAAAAAGCGACCAAATGG |
| 9. OligosP1+P2sense | TAAATCGCATCGTGACCGGTCATAAAAAGACACGTGTCGTCATCGCTATCCAAATAATT GACAAACTCGAAGCCGCCAAGGG |
| 10. OligosP1+P2antisense | CCCTTGGCGGCTTCGAGTTTGTCAATTATTTGGATAGCGATGACGACACGTGTCTTTTTA TGACCGGTCACGATGCGATTTA |
| 11. OligosM1+M2sense | TAAATCGCATCGTGACCAATAATAAAAAGACACGTGTCGTCATCGCTATCCAAATAATTA AAAAACTCGAAGCCGCCAAGGG |
| 12. OligosM1+M2antisense | CCCTTGGCGGCTTCGAGTTTATAAATTATTTGGATAGCGATGACGACACGTGTCTTTTTA TAAAAGGTCACGATGCGATTTA |
| **CHIP** | |
| 13. Fragment A F | AATAGTAAATCGCATCGT |
| 14. Fragment A R | TGGATAGCGATGACGACA |
| 15. Fragment B F | TGTCGTCATCGCTATCCA |
| 16. Fragment BR | CACGATGCGATTTACTA |
